# A dorsal hippocampus-prodynorphinergic dorsolateral septum-to-lateral hypothalamus circuit mediates contextual gating of feeding

**DOI:** 10.1101/2024.08.02.606427

**Authors:** Travis D. Goode, Jason Bondoc Alipio, Antoine Besnard, Devesh Pathak, Michael D. Kritzer-Cheren, Ain Chung, Xin Duan, Amar Sahay

**Affiliations:** Center for Regenerative Medicine, Massachusetts General Hospital, Boston, MA; Harvard Stem Cell Institute, Cambridge, MA; Department of Psychiatry, Massachusetts General Hospital, Harvard Medical School, Boston, MA; BROAD Institute of Harvard and MIT, Cambridge, MA; Department of Ophthalmology, University of California, San Francisco, CA; Department of Physiology, University of California, San Francisco, CA; Kavli Institute for Fundamental Neuroscience, University of California, San Francisco, CA

**Keywords:** context, dorsal hippocampus, dorsolateral septum, eating disorder, feeding, GABA, kappa opioid receptors, lateral hypothalamus, prodynorphin, somatostatin

## Abstract

Adaptive regulation of feeding depends on linkage of internal states and food outcomes with contextual cues. Human brain imaging has identified dysregulation of a hippocampal-lateral hypothalamic area (LHA) network in binge eating, but mechanistic instantiation of underlying cell-types and circuitry is lacking. Here, we identify an evolutionary conserved and discrete Prodynorphin (*Pdyn*)-expressing subpopulation of Somatostatin (*Sst*)-expressing inhibitory neurons in the dorsolateral septum (DLS) that receives primarily dorsal, but not ventral, hippocampal inputs. DLS(*Pdyn*) neurons inhibit LHA GABAergic neurons and confer context- and internal state-dependent calibration of feeding. Viral deletion of *Pdyn* in the DLS mimicked effects seen with optogenetic silencing of DLS *Pdyn* INs, suggesting a potential role for DYNORPHIN-KAPPA OPIOID RECEPTOR signaling in contextual regulation of food-seeking. Together, our findings illustrate how the dorsal hippocampus has evolved to recruit an ancient LHA feeding circuit module through *Pdyn* DLS inhibitory neurons to link contextual information with regulation of food consumption.

**HIGHLIGHTS:**

- DLS(*Pdyn*) neurons receive dense input from the dorsal but not ventral hippocampus
- DLS(*Pdyn*) neurons inhibit GABAergic neurons in the LHA
- Silencing dorsal hippocampus-DLS(*Pdyn*)-LHA circuit nodes abolishes context-conditioned feeding
- *Pdyn* in the DLS is necessary for context-conditioned feeding

## INTRODUCTION

Eating is a necessary and complex series of ingestive behaviors that is heavily sculpted by context. Calibration of feeding is guided by the interoceptive cues from physiological signals that support life and homeostasis^1–3^, associations of hedonic and rewarding experiences that coincide with food consumption^4,5^, and by the linkage of these outcomes to the socioenvironmental cues and exteroceptive contexts in which feeding behaviors are learned and occur^6–11^. Failures to calibrate motivated behaviors, such as food reward-seeking, to these interoceptive and exteroceptive-specific circumstances are core issues for those with eating disorders^12–15^, whose treatment options remain costly, limited, and often only temporarily effective. Patients with an eating disorder may exhibit lengthy and dangerous bouts of food avoidance across contexts (such as with anorexia nervosa or avoidant restrictive food intake disorder) and/or loss of control and overconsumption (as with bulimia nervosa or binge-eating disorder). Illuminating the neurobiology of cellular substrates within a circuit framework that support contextual gating of feeding will edify development of novel therapeutics for eating disorders and for other unwanted or unhealthy eating behaviors.

The dorsal (D) and ventral (V) hippocampi (HPC) make differential contributions to the encoding of contextual information and goals, respectively^16–19^. By encoding details of contexts or environments, the dorsal hippocampus (DHPC) supports spatial and contextual calibration of food-related appetitive behaviors^20–23^. Likewise, the hypothalamus and its many subregions have a deep and rich history of study for their regulation of internal states and learning-dependent changes that contribute to food-seeking and consumption^2,24,25^. Emerging evidence suggests that dysfunction in DHPC-hypothalamic circuits may play a role in disordered eating: functional neuroimaging documented dysregulated responding in the human HPC and lateral area of the hypothalamus (LHA) in the presence of food-associated contextual cues in patients with binge-eating disorder^26^. While this same study noted evidence for LHA input to the human HPC^26^, identities of cell-types and pathways that bridge the DHPC to the hypothalamus^27^, or LHA more specifically, for context-dependent calibration of feeding is absent.

The lateral septum (LS) is a network of local and long-range-projecting inhibitory neurons that receives diverse cortical, subcortical, and dense dorsal and ventral HPC inputs across its own dorsal-ventral axes and in turn projects strongly to multiple areas of the hypothalamus^28–35^. The dorsolateral subregion of the lateral septum (DLS) sits at the top of the LS and is well-positioned to regulate context-dependent behaviors, such as feeding, given its considerable DHPC input and robust targeting of hypothalamic regions, including the LHA^36^. A myriad of cell-types exists within the LS^37–43^, distinguished by their input-output connectivity, physiology and neuropeptidergic expression, and this diversity likely contributes to the various behaviors now associated with LS function^44–49^, including cell-type-specific contributions of LS neurons to stress-responding and conditioned fear^36,50–53^, social interactions^54–57^, and drug- and food-seeking or consumption^43,58–65^. A predominant cell-type in the DLS is somatostatin (*Sst*)-expressing inhibitory neurons^36,43,53^. Using longitudinal calcium imaging, our lab has found that different *Sst* subpopulations are recruited in response to task-specific demands^36^. Thus, DLS(*Sst*) cells, much like cortical *Sst* subpopulations^42,66–69^, are highly heterogenous and are likely to express other distinct neuropeptides, exhibit distinct input-output connectivity, and may broadcast DHPC, VHPC, or convergent HPC inputs to specific subcortical targets to calibrate context-dependent feeding.

Here we identified an evolutionary conserved, dorsally restricted subset of DLS(*Sst*) inhibitory neurons that co-express prodynorphin (*Pdyn*). This small population of DLS(*Pdyn*) inhibitory neurons receive extensive DHPC (but not VHPC) inputs and synapse onto inhibitory neurons of the LHA. We found that inhibition of DHPC-DLS or DLS(*Pdyn*)-LHA circuitry, or deletion of *Pdyn* in the DLS, attenuated the expression of context-specific food consumption, thereby defining a cell type and circuit framework for regulation of context-driven feeding.

## RESULTS

### Prodynorphin-expressing cells are a dorsally biased subset of somatostatin-expressing dorsolateral septal cells

Prior work from our lab revealed that subsets of *Sst*-expressing cells in the DLS exhibit context-dependent expression of task-specific neural activity^36^. Little is known regarding the distribution or identities of potential *Sst*-subtypes within the septum. *Sst*-expressing cells, such as in the cortex^42,66–69^, exhibit a wide range of neuropeptides. Sequencing experiments have noted the expression of various neuropeptides in the septum^36–43^, including *Sst*, neurotensin (*Nts*), proenkephalin (*Penk*), neuropeptide Y (*Npy*), and *Pdyn*, but the extent of their overlap and localization within the LS isn’t well-defined. Moreover, while LS(*Nts*) circuits have been linked to a variety of motivated behaviors^63,65,70,71^, little to nothing is known of other LS neuropeptidergic populations, such as *Pdyn*, *Penk*, or *Npy*. Accordingly, we used multiplex fluorescent *in situ* hybridization (RNAscope^72^) to characterize and identify whether *Nts*, *Penk*, *Npy*, and/or *Pdyn* represent overlapping or unique *Sst*-expressing lateral septal cell populations in male C57/BL6J mice (**Figure 1A** and **S1A**).

**Figure 1.**
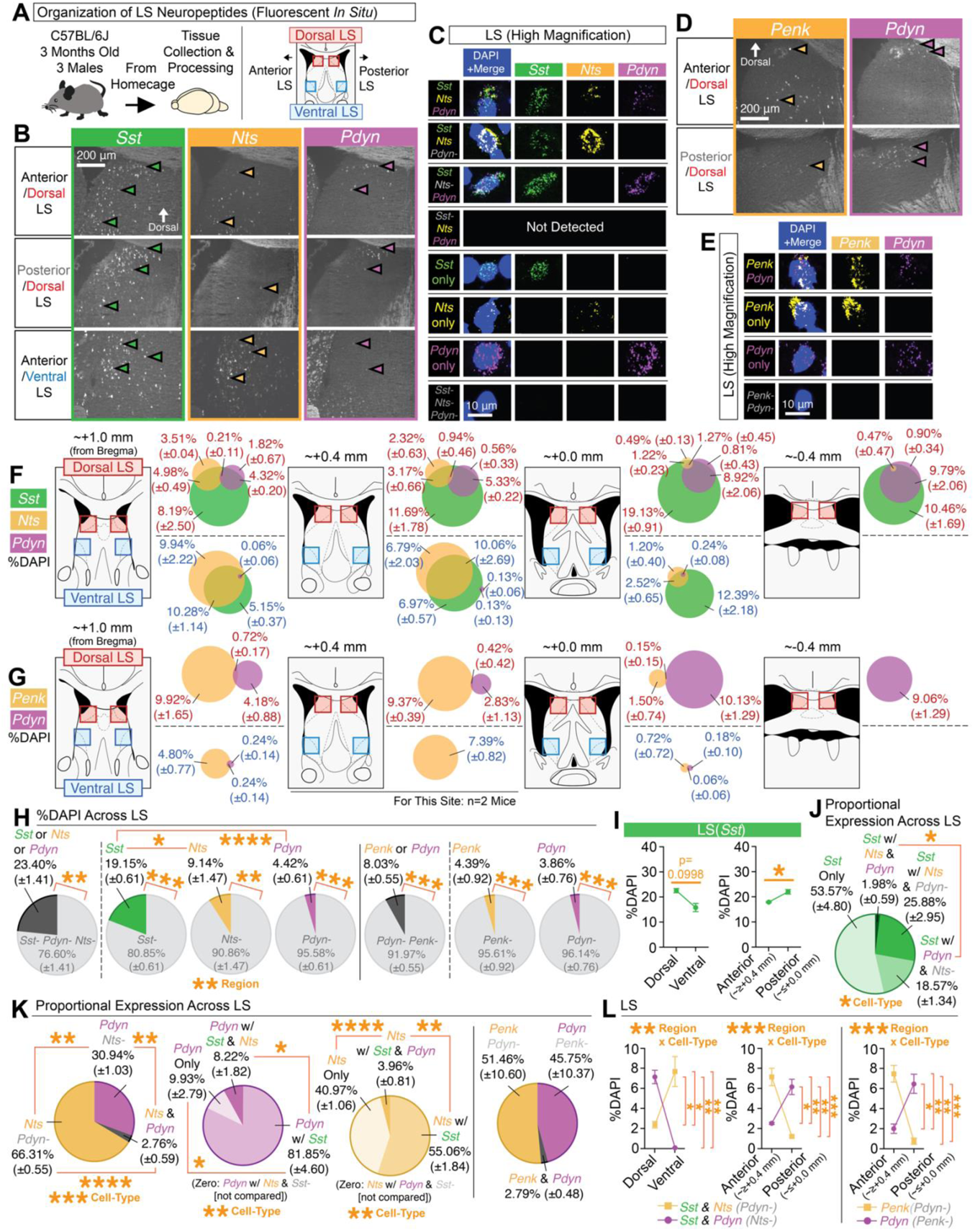
Topographic mapping of neuropeptides reveals *Pdyn* as a distinct dorsal subset of *Sst* cells. **(A)** Multiplex fluorescent *in situ* hybridization was used to map the neuropeptides *Sst*, *Nts*, *Penk*, and *Pdyn* in the LS across its dorsal-ventral and anterior-posterior regions. **(B)** Representative coronal images of *in situs* for *Sst*, *Nts*, and *Pdyn* at different regions of the LS. **(C)** High magnification representative images of individual cells for each of the different cell-types detected for *Sst*, *Nts*, and/or *Pdyn*. **(D)** Representative coronal images of *in situs* for *Penk* and *Pdyn* at different regions of the LS. **(E)** High magnification representative images of individual cells for each of the different cell-types detected for *Penk* and/or *Pdyn*. **(F)** Mouse atlas images and venn diagrams [%DAPI (±SEM)] depicting the extent of overlap of *Sst*, *Nts*, and/or *Pdyn* at each quantified region in the LS. **(G)** Mouse atlas images and venn diagrams [%DAPI (±SEM)] depicting the extent of overlap of *Penk* and/or *Pdyn* at each quantified region in the LS. **(H)** Average expression (%DAPI) of *Sst*, *Nts*, *Pdyn*, and/or *Penk* across all quantified regions of the LS. Main effect of cell-type (ANOVA; significant Tukey’s post-hocs) for comparisons of *Sst*-positive, *Nts*-positive, and *Pdyn*-positive cells. Significant paired t-tests denote comparisons of positive vs. negative expression for each cell-type. **(I)** Average expression (%DAPI) of *Sst* in the dorsal vs. ventral LS and anterior vs. posterior LS (significant paired t-test). **(J)** The average proportion (derived from %DAPI) of each subtype for all *Sst* cells across all quantified regions of the LS (ANOVA: main effect of cell-type; significant Bonferroni post-hoc). **(K)** The average proportion (derived from %DAPI) of each subtype across all quantified LS regions for all *Nts* and/or *Pdyn* cells (ANOVA: main effect of cell-type; significant Tukey’s post-hocs), all *Pdyn* cell-types (ANOVA: main effect of cell-type; significant Tukey’s post-hocs), all *Sst*, *Nts*, and/or *Pdyn* cells (ANOVA: main effect of cell-type; significant Tukey’s post-hocs), and all *Penk* and/or *Pdyn* cells. **(L)** Comparisons of the average expression (%DAPI) of *Sst-Nts* and *Sst-Pdyn* cells in the dorsal vs. ventral and anterior vs. posterior regions of the LS, and Penk or Pdyn cells in the anterior vs. posterior regions of the LS (ANOVAs: significant interactions of region x cell-type; significant Bonferroni post-hocs). For the entire figure, all data are shown as mean (±SEM), and for all statistics: *=p<0.05; **p<0.005, ***p<0.0005; ****p<0.00005.

Representative images of dorsal and ventral LS *in situs* for *Sst*, *Nts*, and *Pdyn* are shown in Figure 1B, with the expression of these genes in individual cells and their combinations shown in **Figure 1C**. The only cell-type we did not observe in the DLS or VLS was *Sst*-negative cells that were positive for both *Nts* and *Pdyn*. Separate LS *in situs* for *Pdyn* and *Penk* are shown in **Figure 1D** (individual cells shown in Figure 1E). We quantified the extent of overlap of all of these neuropeptides as a percentage of the number of 4’,6-diamidino-2-phenylindole (DAPI)-positive cells across multiple sites in the LS (**Figure 1F-1E**). Across all quantified regions, *Sst* was the most abundant cell-type, followed by *Nts*, with similar numbers observed for *Penk* and *Pdyn* (**Figure 1H**). Sst cells were most abundant in the dorsal and posterior LS (**Figure 1I**; also, see **Figure S1F**). Within *Sst*-expressing cells across the DLS and VLS, about half (∼54%) were negative for Pdyn and Nts (**Figure 1J**). Only a very small portion (∼2%) of *Sst*-positive cells expressed both *Nts* and *Pdyn* (**Figure 1J**). A moderate portion of *Sst* cells expressed *Nts* without *Pdyn* (∼26%) and likewise for *Sst* cells expressing *Pdyn* without *Nts* (∼19%; **Figure 1J**). We observed very little overlap of *Nts* and *Pdyn* cells (∼3%) or *Penk* and *Pdyn* cells (∼3%) in the LS (**Figure 1K**). Additionally, while *Pdyn* cells overlapped with *Sst* in ∼90% of its cells, we observed a large portion (∼41%) of *Nts* cells that lacked *Sst* (**Figure 1K**). Moreover, *Pdyn* cells were found to occupy different areas of the LS compared to *Nts* or *Penk*, namely in the more dorsal and posterior regions of the LS (**Figure 1L**).

We also analyzed *Sst*, *Penk*, and *Npy* expression in the LS (**Figure S1A**); representative coronal images of their *in situs* and individual cells are shown in **Figure S1B-S1C**. We did not observe overlap of *Penk* with *Npy* (**Figure S1C**). The extent of overlap across the LS of *Sst*, *Penk*, and *Npy* is shown in **Figure S1D**. We saw similar levels of *Sst* and *Penk* as in the separate *in situs* from **Figure 1**, whereas *Npy* was sparsely expressed in the LS (0.06% of DAPI cells; **Figure S1E**). ∼11% of *Sst*-expressing cells were found to overlap with *Penk* without *Npy*, and while very sparse (∼0.3%), we observed overlap of *Sst* with *Npy* (without *Penk*; **Figure S1G**). Similar to *Nts* (**Figure 1**), we observed a large portion (∼42%) of *Penk* cells that lacked *Sst* (**Figure S1G**). We did observe some (∼33%) *Npy* expression without *Sst*. *Penk* was primarily located the anterior/dorsal region, regardless of whether these cells expressed *Sst* (**Figure S1H**).

Given the high levels of overlap of *Pdyn* with *Sst* (as compared to *Nts*, *Penk*, or *Npy*), and the fact that little to nothing is known regarding the connectivity or role of *Pdyn* cells in the DLS, we further characterized the distribution of *Pdyn* in the LS. To better understand the longitudinal expression of *Pdyn* in the LS, and whether it may be sexually dimorphic, we crossed Pdyn-Cre mice with a developmental reporter line (Ai14) and quantified reporter-labeled (tdTomato) cells across the LS in male and female mice (**Figure S2A**). Representative coronal images showing tdTomato-labeling across the septum are shown in **Figure S2B** (with sites of quantification noted in **Figure S2C**). Similar to our *in situ* data, we observed the densest number of tdTomato-positive cells in the DLS, at levels comparable between males and females (**Figure S2D**). Unlike our *in situ* data, we observed some ventral labeling in the most anterior part of the VLS, suggesting some developmental changes in *Pdyn* expression in the LS, but DLS expression of reporter was roughly three times the amount on average as across the VLS (**Figure S2D**). To observe whether this labeling was static across life, we aged bigenic Pdyn-Cre::Ai14 female mice to 1 year, and performed the same quantifications (**Figure S2E**). None of these quantifications appear to be due to Cre-independent labeling of the reporter line (**Figure S2G**). Thus, *Pdyn* cells appear to be a dorsally biased subset of LS cells that may be similarly distributed in males and females.

A primary question is whether *Pdyn* is conserved in the human LS. To help address this, we accessed the publicly available human *in situ* data of the Human Brain ISH Neurotransmitter Study from the Allen Brain Institute^73^. In manually scanning tissue image sets in these data, we observed at least one specimen with *Pdyn* expression in the LS (**Figure S3**). Screenshots shown are derived from *in situs* of *Pdyn* in human tissue from a 55-year-old male human (control; **Figure S3A-S3B**). The *in situs* include the unilateral LS and neighboring subcortical structures, such as the caudate and accumbens (**Figure S3C-S3D**). In zooming in on the LS, and its dorsal areas in particular (**Figure S3E-S3F**), we observed labeling of a dorsally biased cluster of *Pdyn*-expressing cells in the human LS, suggesting conservation of this cell-type across species.

Together, these data suggested *Pdyn*-expressing cells in the DLS are subpopulation of *Sst*-expressing cells, which are topographically and/or molecularly distinct from other LS neuropeptides, such as *Nts*, *Penk*, or *Npy*. Moreover, DLS(*Pdyn*) cells exhibit similar distributions in both sexes and is conserved in humans.

### Prodynorphinergic neurons in the dorsolateral septum receive dense dorsal hippocampal input

Inputs to DLS(*Pdyn*) neurons are not known. To test for DHPC input, and to begin mapping the afferents to DLS(*Pdyn*) neurons in the brain, we used monosynaptic rabies tracing^74,75^ in male and female mice (bigenic Pdyn-Cre::LSL-TVA), similar to prior reports from our lab^36,50^ (**Figure 2A-2B**). An example starter cell expressing the helper virus (DIO-H2B-GFP-2a-oG) and pseudotyped RG-deleted rabies virus (EnvA-SADΔG-DsRed) the DLS is shown in **Figure 2C**. **Figure 2D** shows a representative coronal image of starter cells and other potential presynaptic partners in the DLS. Across the brain, rabies input mapping revealed several diverse sources of cortical and subcortical input, including dense input from the DHPC (**Figure 2E**). Quantifications of the total number of starter cells in the DLS, the total number of presynaptic cells across the brain, the correlation of these two, and the number of presynaptic cells broken down by region(s) are shown in **Figure 2F**. Normalized as a weighted percentage of each input (totaling 100%), we observed the densest proportion of presynaptic cells in the dorsal hippocampus, namely D/iCA3/2, followed by the LS itself and other dorsal hippocampal regions (e.g., D/iCA1) and so on (**Figure 2G**). In contrast, ventral regions of the hippocampus (VCA1 or VCA3/2) were among regions with the lowest rabies expression. These data indicate that Pdyn-expressing cells are a dorsally biased subpopulation in the DLS that receive considerable DHPC input.

**Figure 2.**
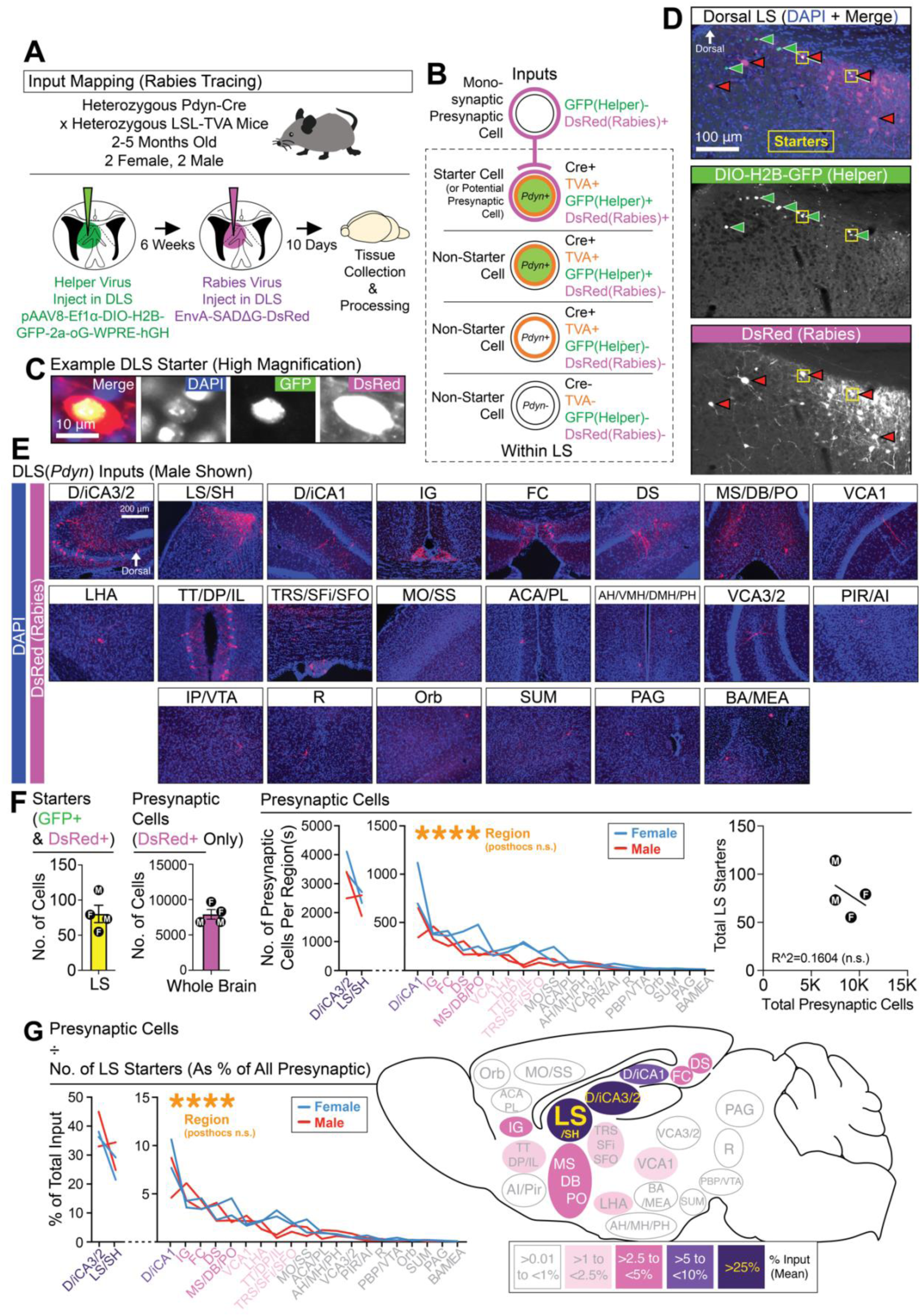
DLS(*Pdyn*) neurons receive dense DHPC input. **(A)** Monosynaptic rabies tracing was used to identify afferents to DLS(*Pdyn*) cells from across the brain. **(B)** A schematic shows the logic used for identifying starter and presynaptic cells. **(C)** High magnification image of a starter cell in the DLS. **(D)** Representative coronal images showing starter cells and monosynaptic labeling in the DLS. **(E)** Representative coronal images of brain-wide inputs to DLS(*Pdyn*) cells. **(F)** The number of starter cells detected in the DLS, the total number and per region of presynaptic inputs (ANOVA; main effect of region), and a plot correlating the number of starters with the total number of presynaptic cells. (**G**) A percentage of total input for each region(s) is calculated and summarized in a sagittal schematic (ANOVA; main effect of region). Outside of the individual data points plotted for each brain region(s), all data in the figure are shown as mean (±SEM); no significant comparisons noted in figure. Abbreviations (see methods for additional details): “D/iCA3/2” (dorsal/intermediate CA3/2 of the dorsal hippocampus), “LS/SH” (lateral septum and/or septohippocampal area within the LS), “D/iCA1” (dorsal/intermediate CA1 of the dorsal hippocampus), “IG” (indusium griseum), “FC” (fasciola cinerea), “DS” (dorsal subiculum), “MS/DB/PO” (medial septum, diagonal band, and/or preoptic area), “VCA1” (ventral CA1), “LHA” (lateral hypothalamic area, which could also include the tuberal area), “TT/DP/IL” (tenia tecta, dorsal peduncular, and/or infralimbic areas), “MO/SS” (motor and/or somatosensory cortices), “ACA/PL” (anterior cingulate and/or prelimbic areas), “AH/VMH/DMH/PH” (anterior hypothalamus, ventromedial hypothalamus, dorsomedial hypothalamus, and/or posterior hypothalamus), “VCA3/2” (ventral CA3/2), “PIR/AI” (piriform area and/or agranular insular area), “IP/VTA” (interpeduncular nucleus and/or ventral tegmental area), “R” (raphe), “ORB” (orbital area), “SUM” (supramammillary nucleus), “PAG” (periaqueductal gray), “BA/MEA” (basal regions of the amygdala and/or medial amygdala).

### Prodynorphinergic dorsolateral septal neurons innervate inhibitory neurons of the lateral hypothalamus

Outputs of DLS(*Pdyn*) neurons are not known. To begin mapping the outputs of DLS(*Pdyn*) neurons, we injected Cre- or Cre- and Flpo-dependent viral vectors into the DLS (**Figure S4A**). We observed the densest fibers in the DLS, the (lateral) diagonal band and preoptic areas (DB/PO), and LHA, with some moderate to sparse expression in the medial septum (MS) and VLS and supramammillary nucleus (SUM; **Figure S4A**). This pattern of expression was similar in males and females and was similar whether we restricted viral expression to depend on both *Pdyn* and *Sst* (**Figure S4A**), reflecting again the high overlap of *Pdyn* with *Sst* in the DLS. In quantifying the density or intensity of DLS(*Pdyn*) fibers in these targets, we measured the fluorescent intensity of virally expressed Synaptophysin-mRuby in these regions (as well as a few regions other efferent regions of the LS that had little or no expression of *Pdyn* fibers; **Figure S4B**). Synaptophysin-mRuby labeling was brightest in the DLS, followed by the DB/PO, LHA, MS, and SUM (with little or no expression in the VLS, or other more medial regions of the hypothalamus) (**Figure S4B**). To support the findings that these output targets are specific to DLS(*Pdyn*) projections (and not contamination from virus spillover into neighboring *Pdyn* structures), we injected Cre-dependent virus into neighboring sites of the DLS in Pdyn-Cre mice, such as anterior cingulate (ACA; dorsal to the DLS), caudate (CPu; lateral to DLS), and dorsal penduncular/infralimbic cortices (DP/IL; anterior to DLS; **Figure S5A-S5C**)—the output patterns of injections in these structures appeared quite distinct from DLS(*Pdyn*) cells. Additionally, and to test for the specificity of many of the viruses used in our experiments, we injected various Cre- or Flpo-dependent viruses into the DLS or LHA of mice that had or lacked Cre or Flpo (respectively)—virus expression was highly faithful to the presence or absence of Cre/Flpo (**Figure S5D-S5G**).

Next, we utilized a virally mediated anterograde labeling strategy^76^ (DIO-mWGA-mCherry) to quantify post-synaptic cells of DLS(*Pdyn*) neurons (**Figure 3A**). We observed a large number of mCherry-positive cells in the DLS (which may or may not be Pdyn-positive) and in the more dorsal areas of the MS (**Figure 3B**)—others’ prior work suggested there may be limited, but not absent, DLS-MS connectivity^77^. The LHA (followed by the SUM) had the highest number of anterogradely labeled cells outside the septum (Figure 3B). Very little cells were observed in the VLS and in the DB/PO (**Figure 3B**; despite the relatively dense fibers we observed in these regions in **Figure S4**). Innervation of the LHA was supported by a large overlap of retrograde tracer (CTb-AF488) from the LHA with tdTomato in the DLS of Pdyn-Cre::Ai14 mice (**Figure S4C**).

**Figure 3.**
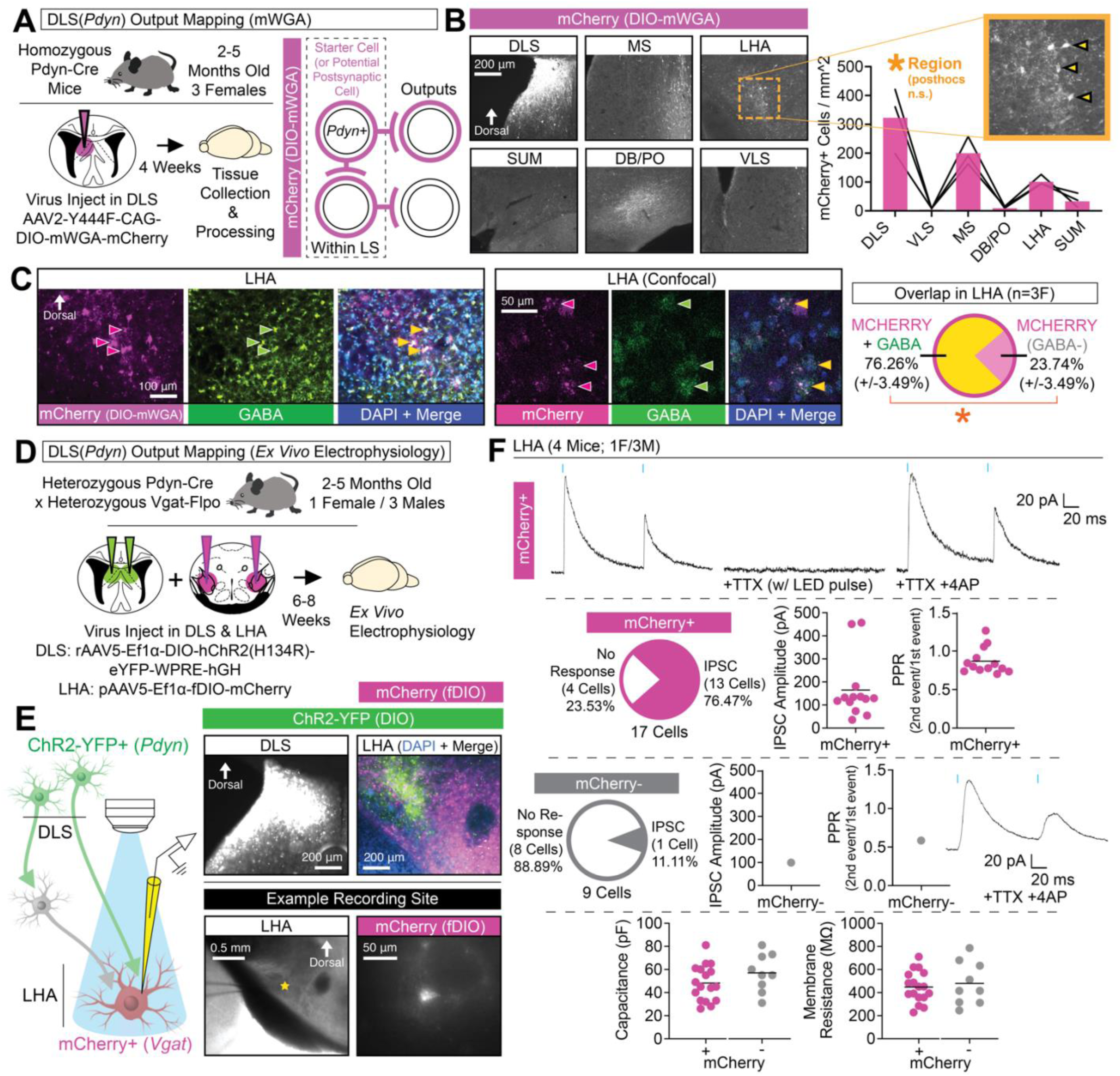
DLS(*Pdyn*) cells project to and inhibit GABAergic cells in the LHA. **(A)** Cell-type-specific and virally mediated anterograde mapping (DIO-mWGA-mCherry) was used to identify outputs of DLS(*Pdyn*) cells in Pdyn-Cre mice. **(B)** Representative coronal images showing DIO-mWGA-mCherry expression in the DLS and output regions, with the quantification of the number of cells per imaged region (ANOVA; main effect of region). **(C)** Representative coronal images (including confocal cross-section images) showing mCherry-expressing cells and immunolabeling for GABA in the LHA, as well as quantification of the percent of overlap across mice in the LHA for mCherry and GABA (significant paired t-test). **(D)** Pdyn-Cre::Vgat-Flpo mice were injected in the DLS with Cre-dependent ChR2-YFP-expressing virus and Flp-dependent mCherry-expressing virus was injecting in the LHA and *ex vivo* electrophysiology was performed. **(E)** Schematic for strategy for ex vivo electrophysiology, with right images showing representative coronal images of ChR2-YFP in the DLS (top left), ChR2-YFP and mCherry in the LHA (top right), and an example recording site (star) and mCherry-positive cell used for patching (bottom right). **(F)** Top: Example traces in a mCherry-positive LHA cell showing inhibitory post-synaptic current (IPSC) following paired pulses of blue light, their loss with TTX, and isolation of monosynaptic responses (+4AP). Top-middle: Quantifications of the number mCherry-positive cells exhibiting light-evoked IPSCs, their amplitude (pA), and PPR. Bottom-middle: Quantifications of the number mCherry-negative cells exhibiting light-evoked IPSCs, their amplitude (pA), and PPR. Bottom: Capacitance and membrane resistance of each recorded cell (w/ and w/o mCherry). No statistical tests were used in (**F)**. For (**C**), the data are shown as mean (±SEM); all other data in the figure are shown as individual datapoints with the mean noted. For all statistics (if applicable): *=p<0.05; **p<0.005, ***p<0.0005; ****p<0.00005.

The LHA is made up of multiple, sometimes non-overlapping cell-types^78,79^ with distinct contributions to feeding—these include vesicular GABA transporter (VGAT)-expressing^80–85,123^ and hypocretin/orexin-expressing cells^86–88^. To begin identifying the efferent targets innervated by DLS(*Pdyn*) neurons in the LHA, we combined our anterograde tracing method with immunofluorescent labeling in the LHA (**Figure 3B**). Immunostaining for GABA in the LHA revealed a large proportion (∼76%) of postsynaptic GABAergic cells (**Figure 3B**). Presumably, anterograde mCherry-expressing cells negative for GABA may consist of LHA’s Vglut2-expressing cell population^78,82,89^. For comparison, we did not observe any orexin-A labeling in anterogradely labeled cells of the LHA (**Figure S4D**).

To confirm monosynaptic inhibitory transmission of DLS(*Pdyn*) cells with GABAergic LHA cells, we used a dual virus (Cre- and Flpo-dependent) strategy in bigenic Pdyn-Cre::Vgat-Flpo mice to optogenetically stimulate DLS(*Pdyn*) terminals in the LHA while simultaneously recording from GABAergic (Vgat-expressing) cells in the LHA using whole-cell *ex vivo* electrophysiology (**Figure 3D-3E**). Representative coronal images of viral labeling in the DLS and LHA, and an example recording site and recorded cell in the LHA, are shown in **Figure 3E**. Representative traces for light-evoked inhibitory postsynaptic currents (IPSCs) following paired pulse in a virus-labeled (Vgat-positive) LHA neuron is shown in **Figure 3F** (top). Synaptic transmission was blocked with tetrodotoxin (TTX), and monosynaptic inhibitory transmission isolated with the addition of 4-aminopyridine (4AP; **Figure 3F**, top). Most of the sampled mCherry-positive cells (17 cells, 4 mice) in the LHA exhibited light-evoked inhibitory transmission (∼77%; **Figure 3F**, middle-top). In contrast, the majority of sampled mCherry-negative cells (9 cells, 4 mice) did not respond to light stimulation (∼89%; **Figure 3F**, middle-bottom). Capacitances and membrane resistances of patched cells were similar between mCherry-positive and mCherry-negative cells (Figure 3F, bottom). We did not observe any light-evoked excitatory postsynaptic currents (EPSCs) for any of the recorded cells (data not shown). Given DLS(*Pdyn*) terminals may release DYNORPHIN, and in preliminary recordings (not shown in figures; 2 mCherry-positive cells from 2 mice, 1M/1F), we also added the KAPPA OPIOID RECEPTOR antagonist, norbinaltorphimine (NorBNI; 1 µM), and observed a slight reduction (mean of ∼-15%) in light-evoked IPSC amplitudes across both cells, suggesting DYNORPHIN release may increase postsynaptic inhibition of LHA(Vgat) cells. Together, these findings are consistent with our histological data that also suggest preferential targeting of GABA-positive cells (**Figure 3C**), and in total, show that inhibitory DLS(*Pdyn*) cells innervate and inhibit GABAergic cells in the LHA.

### Inhibition of DLS(*Pdyn*) neurons disrupts the expression of context-specific conditioned food-seeking

To begin addressing whether DLS(*Pdyn*) neurons may be involved in food consumption, we tested for the overlap of the immediate-early gene, c-Fos, in reporter-labeled cells of bigenic Pdyn-Cre::Ai14 mice (**Figure S6**). Pdyn-Cre::Ai14 mice were assigned to one of three conditions: one in which they would be fasted and given access to a chocolate flavored food in a familiar context, another in which they were fasted but not given access to food in the context, and a final non-fasted group that had access to food in the context (**Figure S6A-S6B**). The mice would then be sacrificed for c-Fos 1 hour after testing for food consumption (or not) in the familiar context. The bodyweight, distance moved, and overall food consumption of the mice are reported in **Figure S6C**; as expected, fasted mice ate the most food. Representative labeling of c-Fos and reporter (tdTomato) in the DLS is shown in **Figure S6D**. Across a similarly sampled proportion of tdTomato-expressing cells in the DLS, we observed significantly more overlap of c-Fos and tdTomato in fasted mice with access to food as compared to without food—non-fasted mice were somewhat in-between the two (**Figure S6**). These data suggest that DLS(*Pdyn*) neurons may be engaged in feeding behaviors.

To better understand how activation of DLS(*Pdyn*) neurons may impact feeding, we next used an optogenetic approach^90^ in freely behaving mice (**Figure S7**). Pdyn-Cre mice were injected with Cre-dependent ChR2-expressing or control virus in the DLS and optic fibers were placed above the DLS (**Figure S7A**). Viral expression and tracts of the optic fibers are shown in **Figure S7B**. Mice then underwent tests for spontaneous feeding in their familiar homecage with concurrent optogenetic stimulation of DLS(*Pdyn*) cells (**Figure S7C**). We observed robust attenuation of food consumption, whether chocolate flavored (**Figure S7D**) or standard chow (outside the homecage, but in a familiar place; **Figure S7E**), with optogenetic stimulation of DLS(*Pdyn*) cells. As reductions in feeding may reflect avoidance behavior^82^, we tested whether optogenetic stimulation triggered avoidance in the real-time place preference (RTPP) assay (**Figure S7G**). We observed some time- and stimulation zone-dependent changes in locomotion (**Figure S7H-S7I**). Moreover, we observed significant avoidance of the stimulation zone in ChR2 mice (**Figure S7J**), suggesting a negative valence^91^ is associated with artificial activation of DLS(*Pdyn*) cells.

We next tried optogenetic inhibition to test for regulation feeding by DLS(*Pdyn*) cells. To begin, mice were given free access to a chocolate flavored food in their homecage (**Figure S8A**) with concurrent optogenetic inhibition of DLS(*Pdyn*) cells (**Figure S8B**). Representative images of viral expression and optic fiber placements across this and other experiments are documented in **Figure S9**. With optogenetic inhibition of DLS(*Pdyn*) cells, we observed an increase in spontaneous consumption of food in non-fasted mice, without a change in the fasted state (**Figure S8C**). These findings are reminiscent of prior work showing increases in food-seeking or consumption in mice over time with inhibition of various cell-types in the LS^58–63^. To determine whether these effects are due to DLS(*Pdyn*)-LHA terminals, in particular, we repeated this test but with optical inhibition of DLS(*Pdyn*) terminals in the LHA (**Figure S8D**). Again, we saw an increase in consumption, this time during the fasted session. While these tests were relatively short, these findings indicate that inhibition of DLS(*Pdyn*) cells, or their terminals in LHA, may promote or augment feeding in a highly familiar setting.

Given the major DHPC input to DLS(*Pdyn*) cells, we hypothesized they may regulate context-dependent forms of consumption. Indeed, rodents can become conditioned to a context to consume more food in that place^6–9^. This process involves repeated pairings of a food reinforcer to a fasted animal in a distinct context; later, the animal is returned to the context in the absence of fasting, and increased consumption is observed, as compared to a separate familiar context that lacks prior reinforcement. Interestingly, others have found increased levels of c-Fos in the lateral septum and LHA (cell-types unknown), following exposure to a food reinforced context^8^. Implementing this behavioral model (**Figure 4C**), we asked whether inhibition of DLS(*Pdyn*) cells (**Figure 4A-4B**) or their terminals in the LHA (**Figure 4E-4F**) regulated the expression of context-dependent feeding. For direct optogenetic inhibition of DLS(*Pdyn*) cells, changes in bodyweight across the experiment, feeding during training, and results during the context tests are shown in **Figure 4D**. Interestingly, rather than broadly increase feeding, we observed consumption that was similarly split across both contexts with DLS(*Pdyn*) inhibition—only controls exhibited context-specific feeding, reflected in the significantly higher discrimination index (**Figure 4D**). Note that this did not interact with whether the A or B context that was reinforced (**Figure 4D**). To determine whether these outcomes are regulated by projections to the LHA, in particular, by DLS(*Pdyn*) cells, we also optogenetically inhibited DLS(*Pdyn*) terminals in the LHA (**Figure 4E-4F**). Bodyweight changes across the experiment and food consumption during training and testing are shown in **Figure 4G**. Optogenetic inhibition of DLS(*Pdyn*) terminals in the LHA eliminated context-induced feeding (**Figure 4G**), in a manner similar to cell-body inhibition of DLS(*Pdyn*) cells (albeit, with somewhat less overall consumption). Overall, these data indicate that DLS(*Pdyn*) cells, and their terminals in the LHA, make critical contributions to the context-specific expression of conditioned feeding.

**Figure 4.**
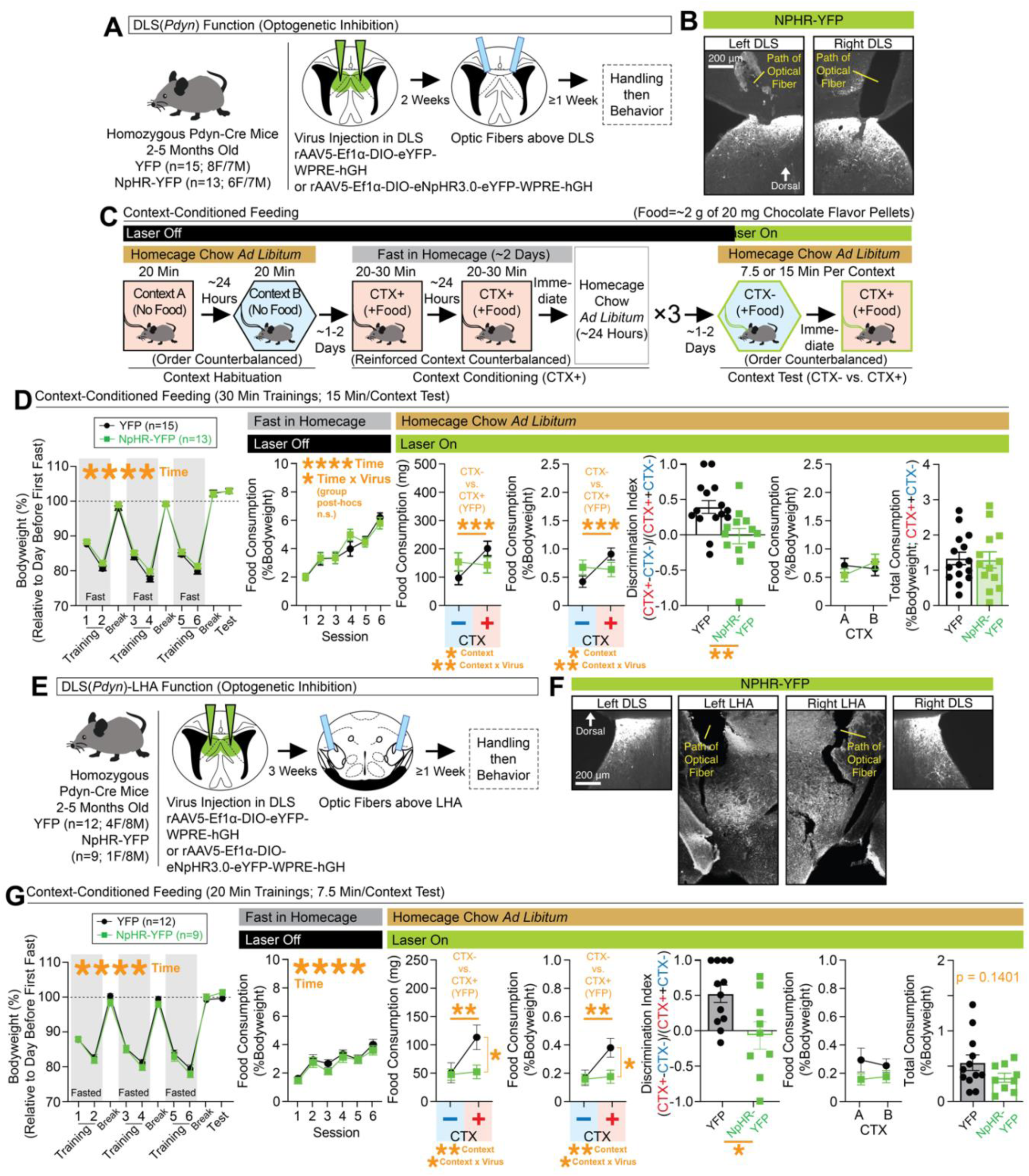
Disrupted expression of context-conditioned food consumption with optogenetic inhibition of DLS(*Pdyn*) cells or their terminals in the LHA. **(A)** Pdyn-Cre mice were injected with Cre-dependent NpHR-expressing or control virus in the DLS and optic fibers were placed above the DLS. **(B)** Representative coronal images with optic fiber tracts and NpHR-YFP-expression in the left/right DLS. **(C)** Behavioral design for context-conditioned feeding. Optogenetic inhibition occurred during the context test phase. **(D)** Leftmost graph depicts bodyweight (%) across training and testing (ANOVA: main effect of time). The next graph depicts food consumption (%Bodyweight) at each training session (ANOVA: main effect of time). At test, food consumption (mg and %Bodyweight) is plotted for the nonreinforced (CTX-) and reinforced (CTX+) contexts (for both mg and %Bodyweight, ANOVAs: main effects of context and context x virus interactions, significant Bonferroni post-hocs for comparing CTX- vs. CTX+ in controls). A discrimination index was generated based on consumption at test (%Bodyweight; significant unpaired t-test). Final two graphs plot consumption (%Bodyweight) across contexts A and B at test (whether CTX+ or CTX-) and total consumption (%Bodyweight) for both contexts at test. **(E)** Pdyn-Cre mice were injected with NpHR-expressing or control virus in the DLS and optic fibers were placed above the LHA. **(F)** Representative coronal images with NpHR-YFP expression in the left/right DLS and LHA and optic fiber tracts above the LHA. **(G)** Leftmost graph depicts changes in bodyweight (%) across training and testing (ANOVA: main effect of time). The next graph depicts food consumption (%Bodyweight) at each training session (ANOVA: main effect of time). At test, food consumption (mg and %Bodyweight) is plotted for the nonreinforced (CTX-) and reinforced (CTX+) contexts (for both mg and %Bodyweight, ANOVAs: main effects of context and context x virus interactions, significant Bonferroni post-hocs for comparing CTX- vs. CTX+ in controls). A discrimination index was generated based on consumption at test (from %Bodyweight; significant unpaired t-test). Final two graphs plot consumption (%Bodyweight) across contexts A and B at test (whether CTX+ or CTX-) and total consumption (%Bodyweight; unpaired t-test shown) for both contexts at test. For the entire figure, all data are shown as mean (±SEM), and for all statistics: *=p<0.05; **p<0.005, ***p<0.0005; ****p<0.00005.

### Attenuation of context-dependent feeding by deletion of *Pdyn* in the septum

*Pdyn* encodes the precursor protein, PRODYNORPHIN (sometimes referred to as PROENKEPHELIN-B), which can be cleaved by PROPROTEIN CONVERTASE 2 (PC2) to yield bioactive forms of opioid neuropeptides, including DYNORPHIN-A, DYNORPHIN-B, and α-NEOENDORPHIN^92^. DYNROPHINS exhibit strong affinity to KAPPA OPIOID RECEPTORS (KORs), such as OPIOID KAPPA RECEPTOR 1 (OPRK1), in the brain^93,94^. In other regions, such as in the nucleus accumbens, DYNORPHIN plays major roles in reward processing^95^. The functional role of *Pdyn* in DLS(*Pdyn*) neurons is not known. Given the effects we observed with inhibition of DLS(*Pdyn*) circuitry, we tested for changes in context-dependent consumption in mice that had conditional knockout of *Pdyn*^96^ in the DLS (**Figure 5**).

**Figure 5.**
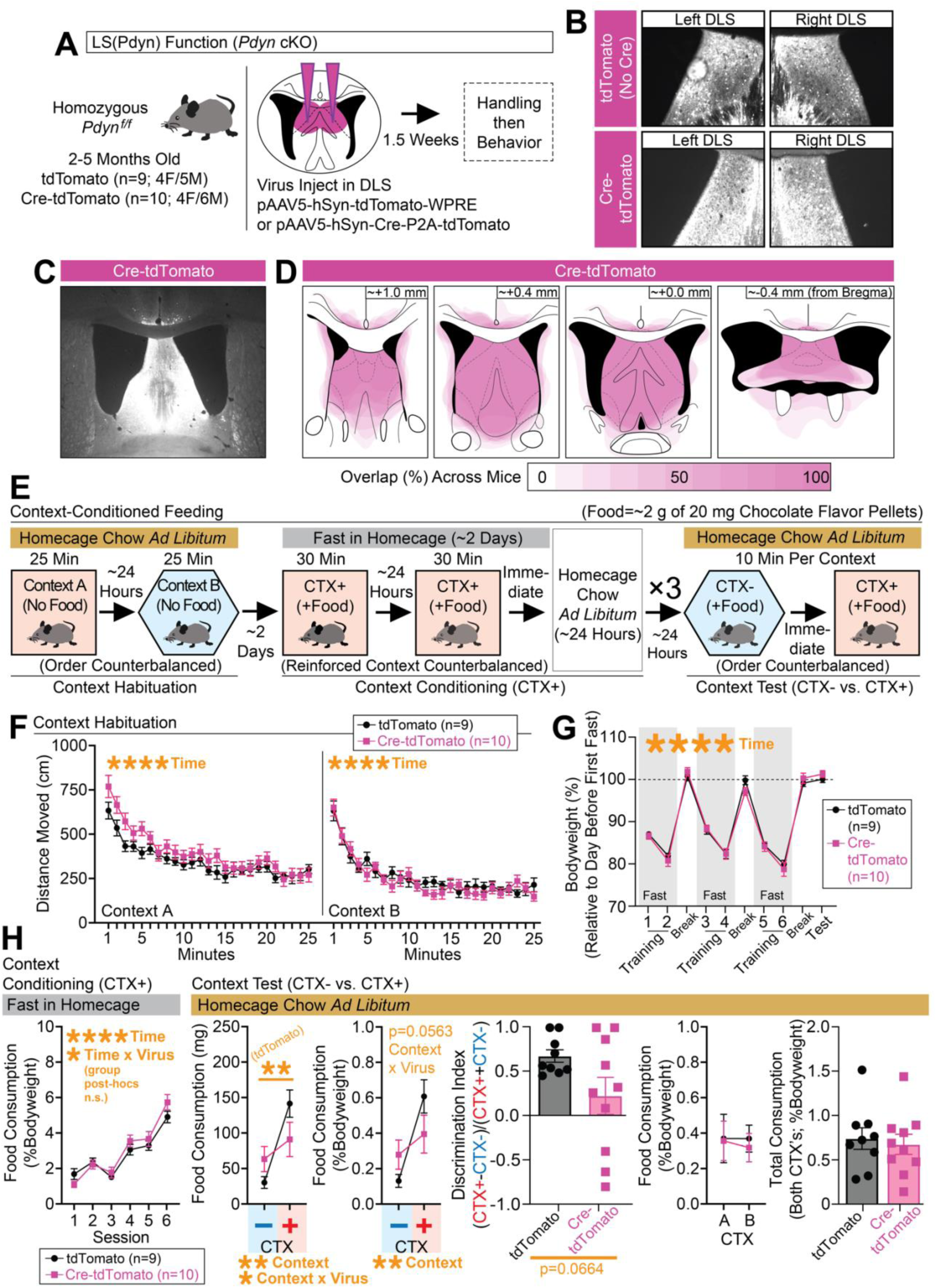
Deletion of *Pdyn* in the DLS alters context-conditioned food consumption. **(A)** *Pdyn^f/f^* mice were injected with Cre-expressing or control virus in the DLS. **(B)** Representative coronal images with Cre-mCherry or mCherry expression in the left/right DLS. **(C)** Larger representative coronal image showing Cre-mCherry expression in the septum. **(D)** Spread of Cre-mCherry virus was documented for each mouse. **(E)** Schematic showing the behavioral design for context-conditioned feeding. **(F)** Distance moved (cm) in contexts A and B during habituation (separate ANOVAs per context: main effects of time). **(G)** Bodyweight (%) across training and testing (ANOVA: main effect of time). **(H)** Leftmost graph depicts food consumption (%Bodyweight) at each training session (ANOVA: main effect of time). At test, food consumption (mg and %Bodyweight) is plotted for the nonreinforced (CTX-) and reinforced (CTX+) contexts (for mg, ANOVA: main effect of context and context x virus interactions, significant Bonferroni post-hocs for comparing CTX- vs. CTX+ in controls; for %Bodyweight, ANOVA: main effect of context, p=0.0563 for interaction). A discrimination index was generated based on consumption at test (%Bodyweight; unpaired t-test shown). Final two graphs plot consumption (%Bodyweight) across contexts A and B at test (whether CTX+ or CTX-) and total consumption (%Bodyweight) for both contexts at test. For the entire figure, all data are shown as mean (±SEM), and for all statistics: *=p<0.05; **p<0.005, ***p<0.0005; ****p<0.00005.

To accomplish this, we injected *Pdyn^f/f^* mice with Cre-tdTomato-expressing or control virus into the DLS (**Figure 5A**) and tested mice in the context-conditioned feeding task (**Figure 5E**). All mice exhibited expression in the DLS (**Figure 5B-5C**), but we acknowledge some spread (although inconsistent) of Cre-mCherry into neighboring structures (documented in **Figure 5D**). Since mice were injected ahead of the start of the behavior and likely had *Pdyn* knocked out in the DLS at the onset of training, we tested for any locomotor changes of the mice during the habituation phase in the two contexts (A and B) that are used for the context-dependent task (**Figure 5F**), but we did not see any significant differences between the groups during this phase. We also tracked bodyweight changes across the training and test phases of the experiment, but did not observe any differences between groups (**Figure 5G**). By the end of training, both groups exhibited similarly high levels of feeding in the reinforced context (**Figure 5H**). At the time of testing, controls exhibited robust context-dependent feeding, but this effect was blunted in Cre-mCherry (**Figure 5H**), albeit with a trending but nonsignificant result when feeding was compared as a discrimination index (**Figure 5H**). As with the other optogenetic inhibition experiments, these effects did not interact with whether the reinforced context was A or B, and the overall consumption across contexts at test was similar (**Figure 5H**). Taken together, these data suggest that *Pdyn* in the DLS is important for the learning and/or expression of context-dependent feeding, findings that mirror the effects seen with inhibition of DLS(*Pdyn*) neurons during expression of context-dependent feeding.

### Context-dependent conditioned feeding depends on dorsal hippocampal projections to the dorsolateral septum, where DLS(*Pdyn*) neurons reside

Given the major innervation of DLS(*Pdyn*) neurons by the DHPC, we next tested whether context-dependent feeding depends on DHPC input to the DLS (**Figure 6**). Animals were injected with CaMKIIα-eNpHR3.0-eYFP-expressing or control virus in the DHPC (aimed at CA3/2, the densest input observed in our rabies tracing) and optic fibers were placed above the DLS (optic fiber placements across all DHPC-DLS experiments are shown in **Figure S9A**). Representative coronal images showing expression of NpHR-YFP in DHPC(CA3/2) cells and their terminals with fibers above DLS are shown in **Figure 6B**. A test for free movement in a novel open field did not reveal any major changes in locomotor behavior with DHPC-DLS inhibition (**Figure S10E-S10H**). Optogenetic inhibition of DHPC-DLS projections in a novel homecage reduced some distance traveled (**Figure S10I-S10J**), perhaps by altering other exploratory behaviors (not measured), given the presence of bedding/sawdust in the cage. When sacrificed 1 hr after exploration in the novel homecage, inhibition of DHPC-DLS projections was found to reduce c-Fos expression in the DLS (an effect that correlated with the overall movement of the mice; **Figure S10K**). To begin testing for effects on context-dependent feeding, we tested fasted mice’ latency and overall consumption in a novel arena (**Figure 6C**). Rodents may reduce their feeding in a novel place relative to a familiar one (termed novelty suppressed feeding^97^). Prior work from our lab found that inhibition of DHPC-DLS projections did not alter the latency to begin feeding when animals were in a novel or familiar environment^50^. Likewise, we did not observe any change in the latency to begin feeding in fasted mice in the current study (**Figure 6D**), but we did see more overall feeding when animals were allowed to continue to feed in the novel arena after initiation of feeding (an effect that was not tested in the prior work)—suggesting a role for this circuit in context-dependent feeding behavior. To further test the contribution of this circuit to contextual feeding, and to observe whether the effects seen with inhibition of DLS(*Pdyn*) cells depends on DHPC input to the DLS, we tested these mice in the context-conditioned feeding task (**Figure 6E**). Bodyweight changes across the experiment, as well as escalating consumption across training, are shown in **Figure 6F-6G**. At test, and similar to our DLS(*Pdyn*) manipulations, we observed a loss of context-specific feeding with inhibition of DHPC terminals in the DLS, as reflected in the discrimination index and overall similar levels of feeding for both contexts (**Figure 6H**; effects that did not depend on whether context A or B was trained). This circuit may be biased to context-regulated forms of feeding as, unlike direct manipulations of DLS(*Pdyn*) neurons, we did not observe any changes in short-term spontaneous consumption in a highly familiar homecage (**Figure S8F-S8G**). Together, these data support a role for DHPC-DLS circuitry in context-dependent consumption of food.

**Figure 6.**
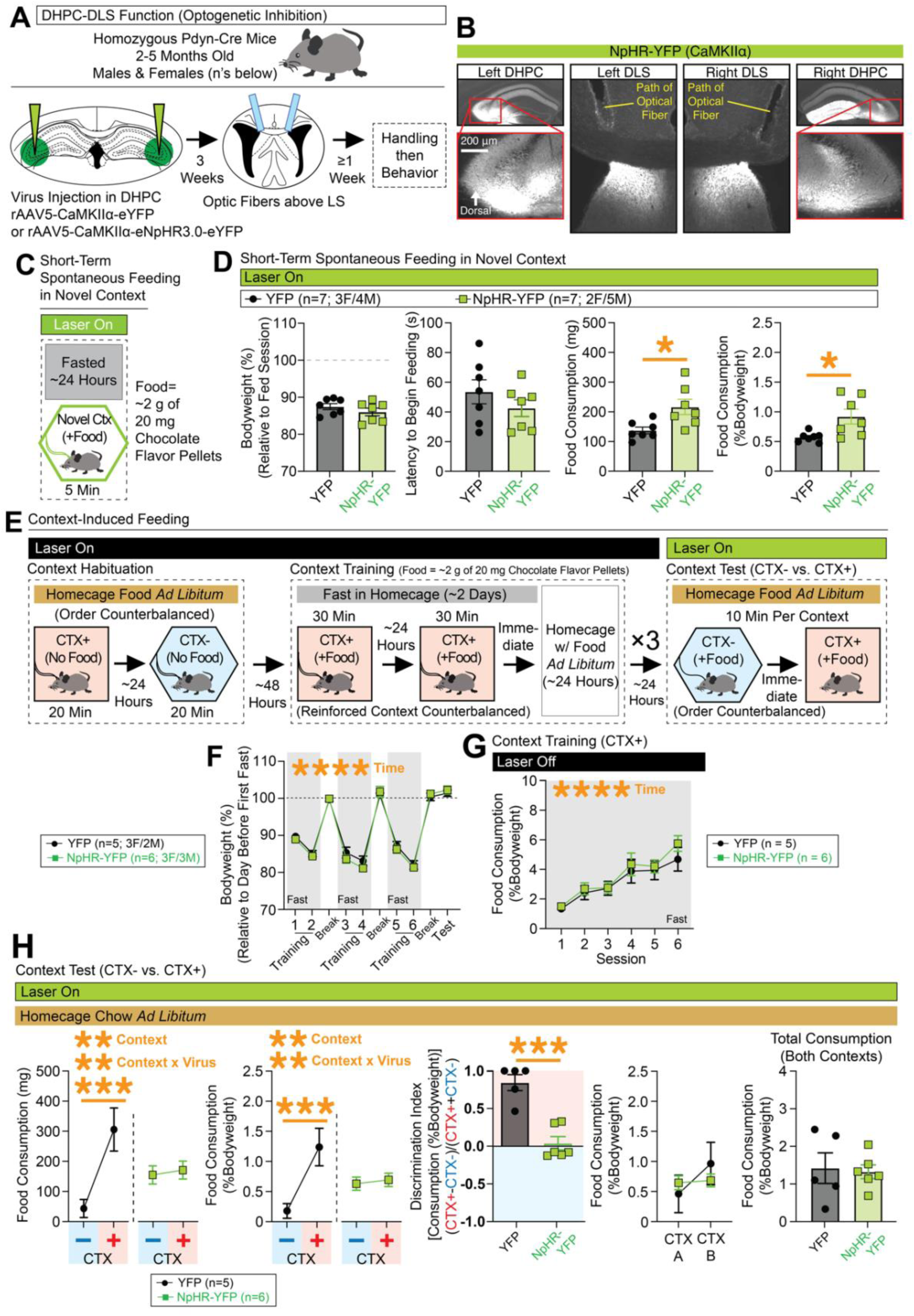
Inhibition of DHPC inputs in the DLS disrupts expression of context-specific expression food consumption. **(A)** Pdyn-Cre mice were injected with NpHR-expressing or control virus in the DHPC and optic fibers were placed above the DLS. **(B)** Representative coronal images with NpHR-YFP expression in the DHPC/DLS and optic tracts above the DLS. **(C)** Behavioral design for testing spontaneous feeding of fasted mice in a novel context. Optogenetic inhibition occurred throughout test. **(D)** Leftmost graph shows bodyweight (%) relative to the day before testing. The next graph shows latency (s) to begin chewing food. The final two graphs show the total amount of food consumed at the end of the test (in mg and %Bodyweight, significant unpaired t-tests for both. **(E)** Behavioral design for context-conditioned feeding. Optogenetic inhibition occurred during the context test phase. **(F)** Bodyweight (%) across training and testing (ANOVA: main effect of time). **(G)** Food consumption (%Bodyweight) at each training session (ANOVA: main effect of time). **(H)** At test, food consumption (mg and %Bodyweight) is plotted for the nonreinforced (CTX-) and reinforced (CTX+) contexts (for both mg and %Bodyweight, ANOVAs: main effects of context and context x virus interactions, significant Bonferroni post-hocs for comparing CTX- vs. CTX+ in controls). A discrimination index was generated based on consumption at test (from %Bodyweight; significant unpaired t-test). Final two graphs plot consumption (%Bodyweight) across contexts A and B at test (whether CTX+ or CTX-) and total consumption (%Bodyweight; unpaired t-test shown) for both contexts at test. For the entire figure, all data are shown as mean (±SEM), and for all statistics: *=p<0.05; **p<0.005, ***p<0.0005; ****p<0.00005.

## DISCUSSION

There is a growing recognition of DHPC contributions to regulation of motivated behavior, however, the neural pathways and cell-types that mediate context-dependent consumption are not known. Here we identify a unique cell-type [DLS(*Pdyn*)] within a multi-node-circuit framework, namely, the DHPC-DLS(*Pdyn*)-LHA pathway, in regulation of context-specific conditioned feeding. Specifically, we found that DLS(*Pdyn*) neurons are a dorsally expressed subset of LS(*Sst*) neurons, which appear molecularly and topographically distinct from other septal neuropeptides, such as *Nts* or *Penk*. DLS(*Pdyn*) neurons receive their densest inputs from the DHPC and project to and inhibit GABAergic LHA populations. These projections calibrate context-dependent behaviors, such that inhibition of DHPC inputs in the DLS or of DLS(*Pdyn*)/DLS(*Pdyn*)-LHA projections altered context-evoked expression of food consumption. DYNORPHIN release may be critical to these processes, as deletion of DLS(*Pdyn*) had a similar impact on the context-dependency of feeding. Activation of DLS(*Pdyn*) neurons altered locomotion and was associated with a negative valence. Additionally, the distribution of DLS(*Pdyn*) neurons was found to be similar in male and female mice, and we observed evidence for its conservation in the human lateral septum. Given that the LS and LHA are ancient circuit modules^98,99^, we speculate that during evolution and emergence of species with hippocampi, these subcortical modules were recruited by the HPC to provide calibration of feeding that is context-specific. In this framework, DHPC outputs, such as from CA3/2, to the DLS may regulate DLS(*Pdyn*)-mediated inhibition of *Vgat*-positive LHA cells to calibrate feeding based on context.

Human imaging suggests that disordered eating may involve functional connections between the DHPC and LHA^26^, processes that may depend on signals bridged by DLS(*Pdyn*) neurons. Inhibition of DHPC-DLS circuitry did not alter pre-learning spontaneous feeding, only context-related feeding on a similar timescale, suggesting a bias of this circuitry in processing or relaying exteroceptive contextual information that is essential for calibrating feeding, or motivation more broadly^100,101^. DLS(*Pdyn*) circuit manipulations affected both spontaneous and conditioned feeding, which may reflect its capacity to integrate this contextual information, as well as its wiring patterns with feeding-regulating cells of the hypothalamus (discussed below). Several studies have identified roles, or potential roles, for distinct cortical^57^, hippocampal^21,27,100–113^, or other subcortical^55,114–117^ inputs to the LS or broader septal areas in regulation of motivated behaviors. Pioneering work using pharmacological dissections have implicated DHPC-LS-LHA orexinergic connections (and to ventral tegmental area, VTA) in the expression of context-dependent forms of drug-seeking^107,108,118,119^. That we do not observe orexinergic projections from DLS(*Pdyn*) neurons (or to VTA) suggests that there may exist multiple parallel pathways for different motivated behaviors in DHPC-LS-LHA circuitry. A lingering question is whether these effects are limited to feeding^121^, or if DLS(*Pdyn*) neurons multiplex to augment other context-dependent behaviors, such as conditioned fear or social behaviors.

DLS(*Pdyn*) neurons are now among several cell-types within the LS that appear to reduce food consumption when stimulated^43,58–64^. Together, these findings suggest that the LS, and perhaps including DLS(*Pdyn*) neurons, may be an important site for therapies targeting overeating and weight loss. Recently, GLP1-R agonists were approved as weight-loss drugs, motivating investigation of circuit underpinnings for their potent anti-satiety effects. Interestingly, GLP1-R expression is observed in the DHPC termination zone of the LS and in the LHA, and stimulation of DLS(*Glp1r*) neurons reduces feeding, suggesting a potential role for these cells within the DHPC-DLS-LHA circuit in contextual regulation of feeding. However, these effects on feeding may come with some caveats, as optogenetic simulation of DLS(*Pdyn*) neurons was also found to induce avoidance, suggesting their activation may trigger negative valence, an effect that could lead to a secondary (not necessarily a primary role) on reduced feeding.

The LHA is a heterogeneous subregion of the hypothalamus, with LHA(*Vgat*) cells exhibiting diverse effects on behavior. In general, ablation or inhibition of LHA(*Vgat*) neurons reduces feeding and may trigger avoidance^80,91,120,122,123^. Accordingly, DLS(*Pdyn*)-dependent changes in feeding or avoidance, such as rapid reductions in homeostatic consumption, may be mediated by their direct inhibition of LHA(*Vgat*) neurons. However, inhibition of LHA-targeting DLS(*Pdyn*) neurons did not increase the overall consumption of mice in the context-dependent task, rather consumption was spread out across the reinforced and nonreinforced contexts, reflecting a loss in context-specificity and not necessarily increases in the expression of feeding itself. Prior work indicates that there are distinct populations in LHA(*Vgat*) neurons that encode or mediate reward associations, or different aspects of feeding, such as approach behaviors preceding food consumption^81^. Thus, one possibility is that manipulations of DLS(*Pdyn*) neurons alters learning-dependent activity within LHA(*Vgat*) cells that is necessary for the expression of context-dependent feeding6, an effect that may be separate from their roles in homeostatic food consumption *per se*. These findings may complement others’ research that implicates hippocampal-septal connections in the expression of food-seeking that is guided by contextual cues^21,103,104,124^, or others’ work that implicates LHA(*Vgat*) neurons in learning-dependent processes for feeding or adaptive behaviors more broadly^6,25,84,125,126^. Other possibilities exist6, however, as DLS(*Pdyn*) neurons exhibit additional connections within the septum and broader hypothalamus (DB/PO, SUM), and we did not see a complete overlap of anterograde labeling of DLS(*Pdyn*) outputs with GABA in the LHA. Perhaps the balance of activity in these and/or other cell-types that receive DLS(*Pdyn*) input are necessary for appropriately matching and expressing the conditioned feeding behavior in a context-specific manner. Future work could determine whether hypothalamic signals that are associated with context-specific consumption are lost with inhibition of DLS(*Pdyn*) inputs.

KOR agonists and antagonists have been shown to have therapeutic potential for a variety of disorders^127–129^, but the cell-type and circuit mechanisms of these effects are still being uncovered. In our study, deletion of *Pdyn* in the septum mirrored the outcomes observed with inhibition of the activity of DLS(*Pdyn*) neurons during the expression context-evoked feeding, suggesting a role for the DYNORPHIN-KAPPA OPIOID RECEPTOR system in DLS circuity, at least at some point along the learning or expression of context-dependent consumption. Given the high levels of consumption by the end of training, these effects may not be due to an inability to engage in or escalate feeding, but rather an inability to attribute this behavior to the appropriate context (either poor contextual learning or discrimination). It is yet unknown what target(s) of DLS(*Pdyn*) neurons are modulated by DYNORPHIN release. One possibility is the LHA itself as sequencing data indicates that various LHA cell populations, including its inhibitory cell populations, express KORs^117,130^. Indeed, and while preliminary, we observed a slight reduction in amplitude of DLS(*Pdyn*)-LHA(*Vgat*)-evoked IPSCs following treatment with the KOR antagonist, NorBNI (see Results section). It should be noted that KOR agonism is sometimes associated with the onset of dysphoria, representing a challenge in the development of KOR therapeutics^131^. As noted above, artificial stimulation of DLS(*Pdyn*) was associated with a negative valence. While it is not yet known whether optogenetic stimulation of DLS(*Pdyn*) neurons is sufficient for DYNORPHIN-release or KOR activation, these findings may relate or contribute to KOR-mediated dysphoric symptoms. Therapies involving KOR agonism or antagonism may consider their effects on DLS(*Pdyn*) circuitry, especially as it may relate to changes in eating behaviors.

In conclusion, alterations in *Pdyn* regulation or dysfunctions in dorsal hippocampus to dorsolateral septum pathway or of dorsolateral septal prodynorphinergic cells to the lateral hypothalamus may result in “circuitopathies” underlying abnormal contextual feeding, and thereby may be novel cellular and circuit substrates for disordered eating or obesity.

## METHODS

### Animal care, subjects, and genotyping

All experiments were conducted in accordance with approved procedures by the Institutional Animal Care and Use Committees (IACUC) at Massachusetts General Hospital and following NIH guidelines (IACUC Protocol # 2011N000084). Mice were housed in a climate-controlled vivarium (22-24 °C) on a reverse day/night cycle with lights on at 7 a.m. and off at 7 p.m. Mice were housed in homecages that included bedding, nesting, and plastic dome, along with *ad libitum* access to water and standard rodent chow (unless stated otherwise as part of experimental conditions). All mice were weaned 3-4 weeks post-birth. Post-weaning, mice were group-housed with 2-4 mice per cage. Breeding mice were housed in their homecages with no more than two adult mice and one litter until weaning. Unless stated otherwise in the figures, mice were 2-5 months of age at the start of any surgery. Ear tagging was used for numerical identification of the mice. Mouse lines and sources are reported in the **Key Resources Table**. All mutant mice underwent standard tail-snip genotyping procedures, using primers and thermocycler protocols corresponding to breeding instructions from their source supplier.

### Transcardiac perfusions and cryosectioning

After completion of experimental procedures, mice were deeply anesthetized with ketamine (105 mg/kg, i.p., Dechra Veterinary Products) and xylazine (7 mg/kg, i.p., Akorn) and transcardially perfused with 20 mL of 1 M phosphate buffer solution (PBS; pH=7.4), followed immediately by ice-cold 4% paraformaldehyde (PFA, in PBS) solution. Brains were extracted and kept in 4% PFA at 4 °C for ∼24 hours before being placed in a 30% sucrose (in PBS) solution at 4 °C until brains sunk in the sucrose solution (∼3 days). Once sunk, brains were placed in small plastic containers filled with freezing embedding medium (Tissue-Tek) and then flash frozen with dry ice. Brains were then stored in a - 80 **°**C freezer until cryosectioning. A Leica cryostat was used for cryosectioning (-20 **°**C). Frozen brains were glued to a mounting stage using the same freezing embedding medium and dry ice and then allowed to acclimate to the cryostat for ∼15 min before sectioning. Unless stated otherwise, 35 μm coronal sections of targeted regions were collected into tissue well-plates containing 0.1% sodium azide in PBS (1 M, pH=7.4) solution. Sectioned tissue was then stored at 4 **°**C until immunohistochemistry or direct wet-mounting onto Superfrost microscope slides (Fisher).

### Multiplex fluorescent *in situ* hybridization

Similar to a prior report from the laboratory^100^, RNAscope^72^ (Advanced Cell Diagnostics; all catalog numbers in this section correspond to this company) was used to visualize mRNA in the lateral septum (**Figure 1** and **Figure S1**). RNAscope was performed on three aged-matched (8 weeks old) naïve male C57BL/6J littermates. The mice were group-housed and allowed to acclimate to the vivarium for 1 wk prior to the following procedures. Animals were sacrificed using rapid decapitation and brains were immediately extracted and flash frozen via 30 sec of submersion in -30 °C isopentane (Sigma-Aldrich). Brains were then kept on crushed dry ice for 1 hr prior to cryosectioning. 10 μm-thick (to minimize cell overlap) coronal sections were collected at -20 °C using a cryostat (Leica; brains were adjusted to the temperature of the cryostat for 30 min prior to sectioning). Serial sections spanning the extent of the LS (approximate locations noted in figures) were placed onto Superfrost microscope slides (Fisher) and kept dry at -20 °C. Sections were stored at -80 °C in a freezer in an air-tight container for 48 hrs. Slides were then removed from the freezer and immediately placed into fresh, ice-cold 4% paraformaldehyde (PFA; in 1 M PBS, pH=7.4) for 15 min. Slides were submerged in fresh 50% EtOH for 5 min at room temperature (RT), followed by fresh 70% EtOH for 5 min (RT), and two 5-min treatments in fresh 100% EtOH (RT). Slides were then allowed to air dry at RT for ∼5 min. Sections were then treated with Protease 3 (Catalog # 322337) for 30 min (RT). Slides were then washed using 1 M PBS, and then submerged in fresh 1 M PBS (RT) for ∼5 min. Slides were then treated with probes from for *Sst* (Mm-Sst; Catalog # 404631), *Pdyn* (Mm-Pdyn-C2; Catalog # 318771-C2), and *Nts* (Mm-Nts-C3; Catalog # 420441-C3), or *Sst* (Mm-Sst; Catalog # 404631), *Npy* (Mm-Npy-C2; Catalog # 313321-C2), and *Penk* (Mm-Penk-C3; Catalog # 318761-C3), or *Pdyn* (Mm-Pdyn-C2; Catalog # 318771-C2) and *Penk* (Mm-Penk-C3; Catalog # 318761-C3). Probes were allowed to hybridize for 2 hrs at 40 °C. Slides were then washed twice (2 min each) with fresh 1× wash buffer (Catalog # 320058). Sections were then treated with AMP1-FL (Catalog # 320852) for 30 min at 40°C, AMP2-FL (Catalog # 320853) for 15 min at 40 °C, AMP3-FL (Catalog # 320854) for 30 min at 40 °C, and AMP4-FL (Alt A) (Catalog # 320855), with washes (as above) after each amplification step. Finally, sections were covered with DAPI Fluoromount-G (SouthernBiotech) and coverslipped. Sections were stored away from light at -20 °C until image collection. These procedures produced three representative sets (*Sst*, *Nts*, and *Pdyn*; *Penk* and *Pdyn*; and, *Sst*, *Npy*, and *Penk*) of LS tissue for three animals (n=3). Using an epifluorescence microscope (Nikon), cells were identified as positively labeled for a given gene based on the detection and localization of aggregate fluorescence above background and attributed to nearest/overlapping DAPI-positive cell. Counts were generated from bilateral images of dorsal and ventral LS cells at four junctures along the rostral-caudal axis (noted in figures). For each mouse, counts were then converted to a percentage of the total number of cells labeled with DAPI in the field of view of the LS, and bilateral percentages were averaged to generate a single value for each DLS region of interest for each probe and for each combination of probes (shown in figures).

### tdTomato expression in Pdyn-Cre::Ai14 mice

Numbers of tdTomato-expressing cells were quantified across the lateral septum of bigenic (heterozygous) Pdyn-Cre::Ai14 littermates, or as compared to age-matched homozygous Ai14 mice lacking Pdyn-Cre (**Figure S2**). Male and female mice were sacrificed at either 2 months or 1 year of age (noted in figure) and coronal sections across the LS were wet-mounted (no immunostain) onto Superfrost microscrope slides (Fisher). Using an epifluorescence microscope (Nikon), two to three coronal images across the bilateral site of interest per mouse were generated of the dorsal and ventral areas of the LS, across its anterior-posterior axes (locations noted in figure). tdTomato-positive cell counts (above background) were manually counted per image and averaged per mouse and normalized to counts/mm^2 by converting the quantified pixel area to mm^2 using ImageJ (NIH). Images were captured with consistent camera exposure and brightness/contrast settings. Wider images of the septum shown in the figure were generated with the same epifluorescence microscope.

### Human lateral septum *in situ* images

Images in **Figure S3** are captured from screenshots of publicly available images from Human Brain ISH Neurotransmitter Study from the Allen Brain Institute^73^.

### Viruses

The following viruses were used, as noted in the figures and Key Resources Table: AAV8-Ef1α-DIO-H2B-GFP-2a-oG-WPRE-hGH (1.54×10^13 gc/ml; Gift of Dr. Xiangmin Xu); EnvA-SADΔG-DsRed (3.5×10^7 gc/ml; Gift of Dr. Xiangmin Xu); AAV2-Y444F-CAG-DIO-mWGA-mCherry (2.0×10^13 gc/ml; Gift of Dr. Xin Duan); AAV5-Ef1α-DIO-hChR2(H134R)-eYFP-WPRE-hGH (4.0×10^12 vg/ml; UNC Vector Core); AAV5-Ef1α-fDIO-mCherry (2.3×10^13 gc/ml; Addgene); AAV5-Ef1α-DIO-eYFP-WPRE-hGH (4.2×10^12 vg/ml; UNC Vector Core); AAVdjd-hSyn-CON-FON-eYFP (5.5×10^12 vg/ml; UNC Vector Core); AAVdj-hSyn-FLEX-mGFP-2A-Synaptophysin-mRuby (1.6×10^13 vg/ml; Addgene); AAV5-Ef1α-DIO-eYFP-WPRE-hGH (3.2×10^12 vg/ml; UNC Vector Core); AAV5-CaMKIIα-eNpHR3.0-eYFP (4.9×10^12 vg/ml; UNC Vector Core); AAV-CaMKIIα-eYFP (5.1×10^12 vg/ml; UNC Vector Core); AAV5-hSyn-tdTomato-WPRE (2.3×10^13 gc/ml; Addgene); AAV5-hSyn-Cre-P2A-tdTomato (1.5×10^13 gc/ml; Addgene). All viruses were stored at -80 °C when not in use.

### Intracranial virus/tracer injections and optic fiber surgery

Mice were first deeply anesthetized with ketamine (100 mg/kg, i.p.; Dechra Veterinary Products) and xylazine (10 mg/kg, i.p., Akorn). To minimize discomfort, mice were treated with carprofen (5 mg/kg, i.p., Zoetis) immediately prior to surgery, and again once per day for three consecutive days after surgery. Hair on the head of the mouse was buzzed short with clippers, and povidone-iodine (Avrio Health) was applied to the skin of the head. Lubricant ophthalmic ointment (Akorn) was applied to the eyes. Mice were then placed in a stereotaxic frame (Stoelting). A small incision was made in the skin at the center of the head and the top of the skull was exposed. The left and right sides of the skull, as well as bregma and lambda of the skull, were aligned on an even horizontal plane. Using a small drill (Foredom), small holes were drilled in the skull above the injection sites. For injections of viruses or tracers into the brain, a pulled glass micropipette attached to a programmable nanoliter injector (Nanoject III, Drummond) was used. Thawed virus or tracer (kept at 4 °C immediately prior to use) was mechanically drawn up into a glass micropipette of the Nanoject III just prior to the lowering of the glass micropipette into the targeted site in the brain. The injector was steadily lowered or raise from the target site over the course of 30 sec and left in place for 1 min before starting the injection and remaining in place for 5 min post-injection. Virus/tracer was injected at a rate of 1 nl/sec until the total injection volume was completed. For injections of viruses or tracers, the following coordinates were used for each region; DLS: 0.0 mm anterior/posterior to bregma, ±0.3 mm from the midline, and -1.9 mm from dura; LHA: -1.3 mm posterior to bregma, ±1.2 mm from the midline, and -5.2 mm from dura; DHPC: -2.0 mm posterior to bregma, ±2.5 mm from the midline, and -2.1 mm from dura; ACA: 0.0 mm anterior/posterior to bregma, ±0.4 mm from the midline, and -1.5 mm from dura; CPu: 0.0 mm anterior/posterior to bregma, ±1.5 mm from the midline, and -2.2 mm from dura; IL/DP: +1.8 mm anterior/posterior to bregma, ±0.3 mm from the midline, and -2.5 mm from dura. For injections into DLS, ACA, CPu, or IL/DP, injection volumes of 0.25 μl/hemisphere were used (reduced to 0.15 μl/hemisphere for rabies tracing experiments in DLS). For injections into LHA, injection volumes were 0.3 μl/hemisphere for tracers or 0.5 μl/hemisphere for virus. Injection volumes were 0.3 μl/hemisphere for DHPC injections. All behavioral/electrophysiological experiments involved bilateral injections, whereas circuit mapping experiments typically involved unilateral injections (noted in schematics in figures). After injection(s), the incision site was closed with sutures (Ethicon). After surgery, mice were removed from the stereotactic frame and mice were monitored for recovery from surgery in a separate clean homecage on a heating pad. Once mice were fully awake and ambulatory, they were returned to their original homecage and allowed to recover for at least a week in the vivarium before any behavioral testing (if applicable). One week before starting behavioral procedures, and just for mice undergoing optogenetic behavioral experiments, a second surgery was performed for implanting optical fibers. This involved all of the same anesthetic and preparation procedures described above prior to drilling. Three small holes were drilled in the skull and three small bone anchor screws (BASi) were attached to the skull. Small holes were drilled for placements of the optical fibers. For optical fibers placed above the DLS, the following coordinates were used (angled at 10° towards the midline): 0.0 mm anterior/posterior to bregma, ±1.2 mm from the midline, and -1.7 mm from dura. For optical fibers placed above the LHA, the following coordinates were used: (angled at 15° degrees towards the midline): -1.3 mm posterior to bregma, ±2.5 mm from the midline, and -4.6 mm from dura. Once in place, dental cement (dyed black with ink) was applied to cover the skull and screws and the bottom half of the optical fibers forming a headcap. Once the headcap hardened, the mouse was removed from the frame and underwent the same recovery procedures described above.

### Monosynaptic rabies tracing

Homozygous Pdyn-Cre mice were crossed with homozygous LSL-TVA mice and their bigenic offspring were used for monosynaptic rabies tracing experiments (**Figure 2**). Pdyn-Cre::LSL-TVA mice were housed in their homecages in a room of the vivarium that was maintained under biosafety level (BSL) 2 conditions. Surgery and euthanasia occurred under BSL 2 conditions and with BSL 2 personal protective equipment. After 6 weeks of expression post-injection of helper virus, mice were injected with rabies virus and then sacrificed for tissue collection after 10 days. Starting at ∼+3.0. mm anterior and continuing through to ∼-5.8 mm posterior to bregma, brains were coronally cryosectioned (35 μm sections) and all tissue from across the brain was wet-mounted to microscope slides and covered with DAPI Fluoromount-G (SouthernBiotech) and glass coverslips (no immunostaining). Using an epifluorescence microscope (Nikon), and across all sections, experimenter(s) manually counted the number of starter cells in the LS, defined as overlap between nuclear GFP expression and cytoplasmic tdTomato expression, as well as the number of tdTomato cells in all regions of the brain where cells were detected. Regions of interest and nomenclature were defined using the Allen Brain Atlas for Adult Mouse Brain (Version 2015)^132^. Some regions were pooled together for their counts. Counts were generated for the following regions of interest: “D/iCA3/2” (dorsal/intermediate CA3/2 of the dorsal hippocampus, defined as anterior to ∼-2.9 mm from bregma), “LS/SH” (lateral septum and/or septohippocampal area within the LS), “D/iCA1” (dorsal/intermediate CA1 of the dorsal hippocampus, defined as anterior to ∼-2.9 from bregma), “IG” (indusium griseum), “FC” (fasciola cinerea), “DS” (dorsal subiculum), “MS/DB/PO” (medial septum, diagonal band, and/or preoptic area), “VCA1” (ventral CA1, defined as CA1 cells past ∼-2.8 mm from bregma), “LHA” (lateral hypothalamic area, which could also include the tuberal area), “TT/DP/IL” (tenia tecta, dorsal peduncular, and/or infralimbic areas), “MO/SS” (motor and/or somatosensory cortices), “ACA/PL” (anterior cingulate and/or prelimbic areas), “AH/VMH/DMH/PH” (anterior hypothalamus, ventromedial hypothalamus, dorsomedial hypothalamus, and/or posterior hypothalamus), “VCA3/2” (ventral CA3/2, from at or posterior to ∼-2.9 mm from bregma), “PIR/AI” (piriform area and/or agranular insular area), “IP/VTA” (interpeduncular nucleus and/or ventral tegmental area), “R” (raphe), “ORB” (orbital area), “SUM” (supramammillary nucleus), “PAG” (periaqueductal gray), “BA/MEA” (basal regions of the amygdala and/or medial amygdala). “% of Total Input” was generated by the following equation: (the number of cells in region of interest ÷ the number of starter cells in the LS) ÷ the total number of presynaptic cells × 100.

### Immunohistochemistry

Unless stated otherwise, visualization of virally expressed fluorophores was enhanced using goat anti-GFP (Novus, 1:500; for YFP-expressing viruses) or rabbit anti-RFP (Rockland, 1:500; for tdTomato-, mCherry- or mRuby-expressing viruses). For GABA and Orexin-A immunostaining, guinea pig anti-GABA (Millipore, 1:400) and mouse anti-Orexin-A (Angio-Proteomie, 1:500) was used, respectively. Rabbit anti-c-Fos (Synaptic Systems, 1:5000) was also used. Secondary antibodies (1:500 for each) included donkey anti-rabbit Cy3 (Jackson ImmunoResearch) for anti-RFP, donkey anti-goat AF488 (Jackson ImmunoResearch) for anti-GFP, donkey anti-guinea pig AF488 (Jackson ImmunoResearch) for anti-GABA, donkey anti-mouse AF488 (Jackson ImmunoResearch) for anti-Orexin-A, and donkey anti-rabbit Cy3 (Jackson ImmunoResearch) for anti-c-Fos. All immunostaining procedures occurred at room temperature aside from incubation of tissue in the primary antibody(s) solution, which occurred at 4 **°**C. All steps occurred with tissue placed on a gentle shaker. Free-floating tissue was transferred between solutions using permeable meshed well inserts (Corning), except for when the tissue was placed in primary antibody solution, as this step and all subsequent steps involved manual transfer of tissue using bent Pasteur pipettes (Sigma-Aldrich). Staining procedures occurred as follows: tissue was washed three times (10 min each) with 1 M PBS (pH=7.4), then placed in 0.01% Triton-X in 1 M PBS for 15 min. The tissue was then washed three more times (10 min each) with 1 M PBS, then placed in blocking solution [10% normal donkey serum (Jackson ImmunoResearch) in 1 M PBS] for 2 hours. Tissue was then washed again (same as before) and then placed in primary overnight (∼12-16 hours). After another set of washings (same as before), the tissue was placed in 1 M PBS containing the respective secondary antibodies for 1.5 hours. Tissue then underwent one final set of washings before being mounted onto Superfrost microscope slides (Fisher) using a paintbrush and 1 M PBS. At the conclusion of staining, the tissue was allowed to dry on the slides at 4 **°**C and then briefly re-wet with 1 M PBS before being covered with DAPI Fluoromount-G (SouthernBiotech) and coverslipped. Antibody concentrations and immunostaining procedures are based on prior work for our lab ^36,50,133,134^ and others ^135^.

### Anterograde virus tracing

AAV2-Y444F-CAG-DIO-mWGA-mCherry was used as an anterograde viral tracer^76^ and injected into the DLS of Pdyn-Cre mice (n’s/sex noted in figure) to map DLS(*Pdyn*) outputs. 4 weeks after injection, mice were sacrificed, perfused, and brain tissue cryosectioned (35 μm coronal sections) and immunostained (anti-RFP). Two to three unilateral images (injection side) per site per mouse of the injection site (DLS, at ∼0.0 mm anterior/posterior to bregma), and potential output regions [MS (towards its more dorsal area, where most cells were found; at ∼0.8 mm anterior to bregma), LHA (at ∼-1.3 mm posterior to bregma), SUM (at ∼-2.8 mm posterior to bregma), DB/PO (at ∼0.5 mm anterior to bregma), and VLS (at ∼0.4 mm anterior to bregma), were imaged with an epifluorescence microscope (Nikon). Numbers of mCherry-positive cells (above background) were manually counted per image and averaged per mouse and normalized to counts/mm^2 by converting the quantified pixel area to mm^2 using ImageJ (NIH). Images were captured with consistent camera exposure and brightness/contrast settings.

### GABA/Orexin-A in the LHA

Confocal z-stack images of mCherry, GABA or Orexin-A, and DAPI expression (∼291 μm x ∼291 μm field of view; ∼40 μm thick z-stacks with ∼1 μm cross-section steps) were generated using a A1R Si confocal laser, a TiE inverted research microscope, and NIS-Elements software (Nikon) for two to three unilateral sections of the DLS (injection side) per mouse. Images were captured with consistent camera exposure and brightness/contrast settings. The numbers of mCherry-expressing cells and their overlap with GABA or Orexin-A (above background) were manually quantified and averaged per mouse. Wider LHA images for the figure were generated using an epifluorescence microscope (Nikon).

### *Ex vivo* electrophysiology

Acute brain slices were prepared and collected using a modified method to improve viability^136,137^. Mice were deeply anesthetized with ketamine (105 mg/kg, i.p., Dechra Veterinary Products) and xylazine (7 mg/kg, i.p., Akorn) then transcardially perfused with ice-cold (4 °C) choline chloride-based artificial cerebrospinal fluid (ACSF) composed of (in mM): 92 choline chloride, 2.5 KCl, 1.25 NaH_2_PO_4_, 30 NaHCO_3_, 20 HEPES, 25 glucose, and 10 MgSO_4_·7H_2_O. Their brains were rapidly extracted following decapitation. Coronal slices (300 µm thick) containing the DLS (for checking viral injections) and LHA (for recordings) were cut in ice-cold (4 °C) choline chloride ACSF using a Leica VT1000 vibratome (Leica Biosystems) and transferred to warm (33 °C) normal ACSF for 30 min. Normal ACSF contained (in mM): 124 NaCl, 2.5 KCl, 1.25 NaH_2_PO_4_, 24 NaHCO_3_, 5 HEPES, 12.5 glucose, 2 MgSO_4_·7H_2_O, 2 CaCl_2_·2H_2_O. All ACSF solutions were adjusted to a pH of 7.4, mOsm of 305, and were continuously saturated with carbogen (95% O_2_ and 95% CO_2_). Slices were allowed to cool to room temperature (20-22 °C) for 1 hour prior to recordings. Whole-cell patch-clamp recordings were amplified, low-pass filtered at 1.8 kHz with a four-pole Bessel filter, and digitized (Multiclamp 700B, Digidata 1550B, Molecular Devices). Slices were placed in a custom-made polytetrafluoroethylene submersion chamber and continually perfused with normal ACSF (>2 mL/min). Neurons were visually identified by infrared differential interference contrast imaging combined with epifluorescence using LED illumination (pE-300 white, CoolLED). Borosilicate patch pipettes had resistances of 4-5 MΩ and were filled with an internal solution containing (in mM): 120 CsMeS, 4 MgCl2, 1 EGTA, 10 HEPES, 5 QX-314, 0.4 Na3GTP, 4 MgATP, 10 phosphocreatine, 2.6 biocytin, pH 7.3, 290 mOsm. Once GΩ seal was obtained, neurons were held in voltage-clamp configuration at -70 mV and the input resistance and capacitance were measured. Series resistance (<30 MΩ) was monitored throughout recordings and recordings were discarded if series resistance changed by >20% from baseline. Excitatory and inhibitory postsynaptic currents (EPSCs and IPSCs) were optically evoked with 1 ms 473 light pulses delivered through the microscope objective. Neurons (mCherry^+^ and mCherry^-^) were identified via optical stimulation through the objective (561 nm). Current responses were recorded at 1.5× threshold, defined as the minimum stimulation intensity required to produce a consistent current response beyond baseline noise. Isolation of EPSC was done by voltage clamp at -70 mV and IPSC at 0 mV. Paired pulse stimulation recordings consisted of 10 sweeps, 1 sweep every 30 s, with a 100 ms interevent interval between pulse stimulation. Paired pulse ratios were analyzed by dividing the amplitude of the second event by the amplitude of the first. Monosynaptic connectivity was assessed by first, confirming the presence of optic evoked IPSCs to paired pulse stimulation. Next, the paired pulse stimulation protocol was conducted after bath application of tetrodotoxin (TTX, 1 µM) to block voltage-gated sodium channels and inhibiting action potential-dependent IPSCs. The stimulation protocol was resumed after bath application of 4-aminopyridine (4AP, 200 µM) to block voltage-gated potassium channels and augment depolarization of ChR2-positive terminals. To assess the influence of kappa opioid receptors on monosynaptic transmission, the paired pulse stimulation protocol was conducted after bath application of norbinaltorphimine (norBNI, 1 µM), a selective kappa opioid receptor antagonist in recordings following TTX and 4AP. 5 min periods between drug application were done prior to the start of each protocol to allow drug diffusion and action. Data acquisition was performed using Clampex and analyzed with Clampfit (Molecular Devices) software.

### Intensity of synaptophysin-mRuby fibers

Homozygous Pdyn-Cre mice were injected with AAVdj-hSyn-FLEX-mGFP-2A-Synaptophysin-mRuby into the DLS and sacrificed after 4 weeks. Coronal sections from across the brain were immunostained (anti-RFP) and two to three unilateral images within the following regions (injection side) were made using an epifluorescence microscope (Nikon; coordinates in mm relative to bregma): DLS (at ∼0.0), DB/PO (at ∼+0.6), LHA (at ∼-1.4), MS (at ∼+0.5), SUM (at ∼+2.8), VLS (at ∼+0.4), AHA/MPO (at ∼-0.5), and VMH (∼-1.5). Images were captured with consistent camera exposure and brightness/contrast settings. Then, images were opened in ImageJ (NIH) and a circle with a diameter of 350 pixels was placed over the middle of the region(s) of interest and the image’ mean intensity or brightness within the circle was measured [in artificial units, ranging from zero (pure black) to 255 (solid white)] using ImageJ’s “Measure” feature and reporting the “mean” value. Mean intensity of each region is then plotted with the mean value of the region of lowest intensity subtracted and shown per mouse.

### Retrograde tracing using recombinant cholera toxin subunit B

To serve as a retrograde neural tracer, 100 μg of recombinant cholera toxin subunit B conjugated to Alexa Fluor 488 (CTb-AF488; ThermoFisher) was dissolved in 500 μl of 1 M PBS (pH=7.4) and stored at -20 °C when not in use. Homozygous Pdyn-Cre and homozygous Ai14 mice were bred together and their bigenic (heterozygous) offspring (Pdyn-Cre::Ai14) were unilaterally injected with 0.3 μl of CTb-AF488 into the LHA. After 1 week, mice were sacrificed, and brain tissue was processed to generate coronal sections (35 μm) of the DLS and LHA (no immunostaining was performed). Confocal z-stack images of tdTomato, CTB-AF488, and DAPI expression (∼291 μm x ∼291 μm field of view; ∼40 μm thick z-stacks with ∼1 μm cross-section steps) were generated using a A1R Si confocal laser, a TiE inverted research microscope, and NIS-Elements software (Nikon) for two to three unilateral sections of the DLS (injection side) per mouse. Images were captured with consistent camera exposure and brightness/contrast settings. The numbers of tdTomato- and CTb-AF488-expressing and the extent of their overlap were then quantified and averaged per mouse. Wider DLS images for the figure were generated using an epifluorescence microscope (Nikon).

### Neighboring *Pdyn* projections

For qualitative comparisons of outputs of neighboring *Pdyn*-expressing regions near the DLS (**Figure S5**), unilateral injections were made of AAV5-Ef1α-DIO-hChR2(H134R)-eYFP-WPRE-hGH into the anterior cingulate area (ACA), caudate putamen (CPu), or dorsal peduncular and/or infralimbic cortices (DP/IL). After 4 weeks, mice were sacrificed, and brain tissue processed. Using an epifluorescence microscope (Nikon), representative coronal images were made of the injection sites, and of the following regions, to match areas where DLS(*Pdyn*) fibers were observed: DLS, VLS/MS, DB/PO, LHA, and SUM. Other regions that had detectable fibers from ACA(*Pdyn*), CPu(*Pdyn*), or DP/IL(*Pdyn*) injections are also shown, including: anterior cingulate and/or prelimbic areas (ACA/PL), prelimbic, infralimbic, and/or dorsal peduncular areas (PL/IL/DP), piriform area and/or agranular insular area (PIR/AI), basolateral and/or central amygdala (BLA/CEA), periaqueductal gray (PAG), ventral and lateral CPu, dorsal and ventral bed nucleus of the stria terminalis (BST), substantia nigra (SN), and nucleus accumbens (NAc). Images were captured with consistent camera exposure and brightness/contrast settings.

### Virus specificity

The following viruses/mice (n’s/sex noted in the figure) were used in tests for virus specificity (**Figure S5**): AAV5-Ef1α-DIO-eYFP-WPRE-hGH was unilaterally injected into the DLS of homozygous Pdyn-Cre or homozygous Sst-Flpo mice; AAVdjd-hSyn-CON-FON-eYFP was unilaterally injected into the DLS of bigenic (heterozygous) Pdyn-Cre::Sst-Flpo or homozygous Sst-Flpo mice; AAVdj-hSyn-FLEX-mGFP-2A-Synaptophysin-mRuby was unilaterally injected into the DLS of of homozygous Pdyn-Cre or homozygous Sst-Flpo mice; and, AAV5-Ef1α-fDIO-mCherry was unilaterally injected into the LHA of bigenic (heterozygous) Pdyn-Cre::Vgat-Flpo or homozygous Pdyn-Cre mice. After 2-4 weeks (noted in the figure), the mice were sacrificed and coronal sections (no immunostaining) of DLS and LHA (relative locations noted in the figure) were imaged for YFP-, mRuby-, or mCherry-expressing cells using an epifluorescence microscope (Nikon). Images were captured with consistent camera exposure and brightness/contrast settings. Cell counts (above background) were manually counted per image and averaged per mouse and normalized to counts/mm^2 by converting the quantified pixel area to mm^2 using ImageJ (NIH).

### Optical fibers and laser delivery

Optical fibers for *in vivo* optogenetics were constructed in the lab. Optical fibers (200 μm core, 0.37 numerical aperture multimode fiber; Thorlabs) were scored and cut with a ruby fiber optic scribe (Thorlabs) and stripped of external coating with a fiber stripping tool (Micro Electronics) and threaded through and glued with epoxy (Thorlabs) in a 230 μm core zirconia ferrule (Precision Fiber Products). 2 or 6 mm of optical fiber was extended out of the bottom of the ferrule for DLS- or LHA-targeting experiments, respectively. The top of the fiber was manually polished with ferrule polishing paper (Thorlabs), and a fiber inspection scope (Thorlabs) was used to ensure smooth polishing of the top of the ferrule. For optogenetic experiments comparing ChR2-expressing viruses to controls, a 100 mW 475 nm blue laser diode was used (OEM Laser Systems). 475 nm laser output was pulsed at 15 Hz with a 20 ms pulse width using an external arbitrary waveform generator (Agilent) attached to the signal port of the laser. For optogenetic experiments comparing NpHR-expressing viruses to controls, a 100 mW 561 nm green/yellow laser diode was used. Laser beam output was collected through a non-contact collimator connected to a 2 m FC optical patch cable (Thorlabs) attached to an FC/PC rotatory optical commutator (Doric). Two 1 m PC optical fiber cables (Doric) extended from the commutator and would be attached to the optic fibers in the headcap of the mice via external zirconia sleeves (Precision Fiber Products), completing the connection of the laser to the optic fibers in the headcap and allowing for free movement of the mice. The output power of the 475 nm and 561 lasers was calibrated to be ∼6-8 mW at the tip of any polished fiber used in the experiments. Laser power was measured using an optical power meter (Thorlabs). When optogenetics were used in behavioral experiments, mice remained in their respective transport container for ∼30 sec after attaching the optic fiber cables, just before being placed in the behavioral arena for testing as described. These stimulation parameters were selected based on prior work from the lab that used these same parameters for stimulation or inhibition of DLS(*Sst*) neurons and DHPC-DLS projections^36^.

### General overview of behavioral procedures

To age-match groups across treatments, experimental mice within each cage were randomly assigned to a viral treatment at surgery, with group assignments balanced equally as best as possible within each cage. During behavioral procedures, experimenter(s) were aware of numerical identification of the mice but remained blinded to the virus or group assignments until after the experiment. Experimenter(s) were not blind to the sex of the mice during behavioral testing or whether food was present or absent. The test order of subjects throughout behavioral procedures was predetermined and pseudorandomized (i.e., the order of testing typically alternated across virus groups). Typically, all male or female mice would complete the task on a given day before switching to other sex. Within a given behavioral task, the order of the mice was kept consistent across days. All behavioral procedures occurred in separate quiet behavior room of the laboratory that had adjustable lighting. For all behavioral procedures, mice were brought to/from the vivarium to the behavior room in their homecages and were allowed to acclimate to the behavior room for 30 or more min prior to starting any behavioral procedures. Behavioral procedures occurred during the day cycle of the mice’ day/night cycle. ∼30 min prior to beginning any behavioral test, animals were placed into a temporary homecage, and returned to their group-housing once all cage-mates completed the day’s tests. This was done to prevent any fighting between animals immediately prior to the start of behavior. Water was provided in the temporary homecage (standard rodent chow was added if mice were not undergoing a fast). All phases of behavioral training were filmed from above in full view of the mouse and arena using a digital USB camera (Microsoft). Lux levels for each arena are reported for each task, as measured using a light meter (Fisher Scientific). Some experiments had multiple cohorts of animals (noted below), which were balanced across viral treatments and sex distributions as best as possible; data shown in figures are collapsed across cohorts (if applicable) for the same experiment. For behavioral experiments using optogenetic inhibition of DLS(*Pdyn*) cells, two cohorts were used: one cohort undergoing just the context-conditioned feeding task, while a second cohort underwent the context-conditioned feeding task followed later by the spontaneous feeding task in the homecage. For behavioral experiments using optogenetic excitation of DLS(*Pdyn*) cells, three cohorts were used: one cohort underwent just the spontaneous feeding task in the homecage, another cohort underwent the real-time place preference task followed by a test for spontaneous feeding of regular chow in a familiar arena, and a third cohort underwent the real-time place preference task followed by the spontaneous feeding task in the homecage. For the behavioral experiment looking at c-Fos expression in Pdyn-Cre::Ai14 mice, a single cohort was used. For behavioral experiments using optogenetic inhibition of DLS(*Pdyn*)-LHA terminals, a single cohort was used for all experiments. For behavioral experiments using conditional deletion of DLS(*Pdyn*), one cohort was used for the context-conditioned feeding task. For behavioral experiments using optogenetic inhibition of DHPC-DLS terminals, two cohorts were used: one cohort underwent novel open field exposure, a test for spontaneous feeding in the homecage, spontaneous feeding in a novel context, and then exposure to a novel homecage. A second cohort underwent novel open field exposure, a test for spontaneous feeding in the homecage, the context-conditioned feeding task, and then exposure to a novel homecage.

### Handling

Mice were handled by the experimenter(s) prior to undergoing any behavioral testing. Mice were brought in their homecages from the colony to the laboratory and allowed to acclimate in their homecages to the behavioral testing room for at least 30 min prior to the start of handling. Handling consisted of the experimenter(s) gently touching and picking up each mouse over the course of ∼30 sec per day for three consecutive days. For mice that were undergoing optogenetic experiments, and in addition to the handling procedures stated above, handling also included attaching the optic fiber cabling to the optic fibers on the headcap of the mouse. Mice were allowed to freely move in a temporary homecage for ∼30 sec before removal of the optic fiber coupling and being returned to their original homecage.

### Food

Unless stated otherwise, Dustless Precision Pellets (20 mg, chocolate flavored, rodent purified diet; Bio-Serv) were used in tests for food consumption. Mice were habituated to the chocolate flavored pellets by spreading ∼2 g of the pellets in their homecages once per day for three consecutive days prior to any tests for feeding behaviors (typically as part of handling procedures). A clean 9 cm wide polystyrene disposable petri dish (VWR) with the lid removed was used as a container to hold and measure the amount of food consumed during tests. Removable mounting putty was attached to the underside of the petri dish and used to adhere the dish to the floor of any testing arena. Food consumption was measured (in g, to the fourth decimal) by weighing the food (∼2 g added for any given test), the petri dish, and putty together on a scale (Mettler Toledo) before and after the task and converted to a difference in mg. To reduce its novelty, a clean, empty petri dish was placed in the homecage of the mice during handling procedures. When standard rodent chow was used in consumption tests, ∼8 g of the chow was placed in the petri dish (the chow was first manually broken into smaller pieces to aid consumption) and measured the same way. Any food that was dropped or carried outside the petri dish by the mouse was added back to the petri dish, and any debris, urine, or feces was remove from the petri dish before being weighed. Mice were weighed (in g, to the first decimal) on a scale (Ohaus) ∼5 min or less before any test in which consumption was compared to their bodyweight. Consumption as a percentage of bodyweight was used to account for differences in weight loss, age, or sex distribution across groups. For example, a heavier male mouse may eat more when fasted, but this consumption may be similarly proportional to its bodyweight as compared to the consumption of a smaller fasted female mouse.

### Motion tracking

For some experiments (**Figure 5**; **Figure S6**; **Figure S7**; **Figure S10**), locomotion (distance moved) and/or location (body centerpoint) data were generated using Ethovision XT Version 15 (Noldus). Video recordings of behavior were uploaded in the software, the borders and size of the arena were noted within each video, and each mouse’s body centerpoint was tracked using built-in autodetection and tracking of the software. Videos and tracking profiles were manually inspected to ensure reliable and consistent tracking of the mice.

### Context-conditioned feeding task

We tested for context-dependent feeding (**Figures 4**, **5**, and **6**) using a modified version of a context-conditioned feeding task used by others in rats and mice^6–9^. Two different contexts were used (light levels were adjusted to 50 lux in the very center of both arenas). Context A consisted of the following features: a 40 cm x 40 cm x 40 cm square plastic arena, with walls and floor covered in gray gaffer tape, and wiped down before/after with 70% EtOH. When present, the food tray would be placed along the midpoint of one wall of the context. Animals were transported to and from the temporary homecage and context A in a gray square plastic container. The second context (B) consisted of the following features: a 40 cm tall white plastic posterboard was wrapped in a circular shape within a 40 cm x 40 cm x 40 cm plastic arena to generate circular arena. Vertical black tape ran down two opposing sides of the circle’s walls. The floor was a perforated metal sheet that had 1 cm wide openings tiled 1 cm apart (edge to edge) throughout the floor, filled in by resting on a piece of opaque plastic. This context was wiped down before/after with 0.1% acetic acid. When present, the food tray would be placed next to one of the black stripes. Animals were transported to and from the temporary homecage and context B in a round white plastic container. The following procedures were used (exact times of each session are noted in the figures): first, to habituate mice to the contexts, mice were exposed to one context per day (pseudorandomized and counterbalanced for the starting context) without food in the contexts, and while they still had access to food in their homecage. A day after the final habituation, context reinforcement training began and consisted of six feeding sessions in one of the contexts (termed “CTX+”, the nonreinforced context is termed “CTX-”). The night before the first session (∼7 pm), homecage food was removed, and on the following two days, the mice were given free access to the chocolate flavored food for 20-30 minutes (noted in figures) in the CTX+ once per day. Immediately after the second training session in the CTX+, mice were returned to their homecage with standard homecage chow present for at least 24 hours. This process was repeated three times for a total of six training sessions, and then mice were maintained with homecage chow in their homecage throughout the rest of the behavior. ∼48-72 hours after the sixth and final training session, mice were tested for context-dependent consumption by comparing consumption in in the reinforced context (CTX+) immediately before or after (order counterbalanced) equivalent exposure to the previously habituated context (CTX-). Bodyweight measurements during “breaks” listed in the figures corresponds to the day before the next phase of the experiment. The equation for the discrimination index is as follows (typically, using %Bodyweigtht consumption): [(Consumption in context CTX+ – Consumption in context CTX-) ÷ (Consumption in context CTX+ + Consumption in context CTX-)]. Mice were placed in the center of the arena whenever first starting in the context.

### c-Fos expression after feeding in Pdyn-Cre::Ai14 mice

Bigenic (heterozygous) Pdyn-Cre::Ai14 mice were tested for food consumption in a habituated context, with or without fasting, and then sacrificed for c-Fos to observe overlap with tdTomato-expressing cells in the DLS (**Figure S6**). Mice were randomly assigned to one of three groups: non-fasted mice with access to food at test in familiar place [“Fed (Food at Test)”], mice that were fasted for ∼24 hours but were exposed to familiar context in the absence of food [“Fasted (No Food at Test)”], or fasted mice with access to food in a familiar place [“Fasted (Food at Test”)]. All mice were first individually exposed to the test context for 20 min prior to testing the following day. Testing consisted of 30 min of exposure to the context again with or without food, and with or without fastings (as noted). All mice were sacrificed and perfused for c-Fos 1 hour after the end of the test. Distance moved of the centerpoint of mouse was tracked using Ethovision XT (Noldus). Confocal z-stack images of tdTomato, c-Fos, and DAPI expression (∼291 μm x ∼291 μm field of view; ∼40 μm thick z-stacks with ∼1 μm cross-section steps) were generated using a A1R Si confocal laser, a TiE inverted research microscope, and NIS-Elements software (Nikon) for two to three z-stacks across the bilateral DLS per mouse. The number of tdTomato-postive, c-Fos-positive, and their overlap (above background) were manually counted and are shown as an average count or percentage per field of view per mouse. Images were captured with consistent camera exposure and brightness/contrast settings.

### Short-term spontaneous feeding in a familiar homecage

Mice were tested for spontaneous feeding in their homecage with and without undergoing a fast of ∼24 hours (**Figure S7** and **Figure S8**). The roof, food tray, and water bottle were removed from the animal’s homecage and a petri dish containing food was placed on the floor of the homecage along a wall opposite the homecage’s water lixit. This homecage was then placed in a larger walled arena to prevent the animal from climbing out of the homecage but allowing for video filming and for the free movement of the optic fiber cables. Light levels were adjusted to 50 lux at the center of the homecage. For homecage feedings, consumption was visually confirmed by experimenter(s) from behind a curtain, as it was not uncommon for non-fasted mice to begin spreading bedding into the petri dish (a primary reason for limiting the sessions to 5 min). Any debris from the homecage was removed from the petri dish before being weighed.

### Short-term spontaneous feeding in a familiar arena (standard rodent chow)

Mice were tested for spontaneous feeding of standard rodent chow in a familiarized arena after being fasted in their homecage for ∼24 hours (**Figure S7**). A familiar arena was used instead of the homecage to avoid the rodent chow (which crumbles easily) from being lost or mixed in the homecage bedding, which could complicate food consumption measurements. The arena was the same arena as used for open field testing (novel for this cohort during the habituation session), but with the food/container being placed in one corner of arena at test (and with reduced lux levels: 50 lux at its center). The mice were transferred to/from the familiar arena in their temporary homecage and gently placed in the center of the arena at the start of the behavior session. Just prior to starting the fast, animals were first habituated to the arena and optic cables (no laser stimulation) for 20 min and allowed free movement. During the test the next day, the laser was turned on immediately when the animal entered the arena and with free access to ∼8 g of rodent chow in the petri dish in a corner of the arena.

### Real-time place preference (RTPP)

RTPP (**Figure S7**) occurred for 15 min in a 60 cm x 30 cm x 30 cm white rectangular plastic arena (Maze Engineers). The midpoint of the arena was divided by two 10 cm wide walls with a center entry point. The light levels in the centers of both sides of the arena were adjusted to 50 lux each. To trigger optogenetic stimulation when the animal was only in one side of the chamber, a digital USB camera (Microsoft) filming from above was connected to a computer running Ethovision XT (Noldus). An external TTL-USB adaptor (Noldus) bridged the laptop and arbitrary waveform generator and laser source. The borders of the arena were established in Ethovision XT (Noldus) and detection of the body centerpoint of the mouse in the stimulation side triggered the immediate activation and pulsing of the laser. The stimulation side was predetermined to counterbalance the stimulation zone across group assignments. To begin, mice were gently placed in the center of the arena between the divider walls.

### Short-term spontaneous feeding in novel arena

Mice were tested for spontaneous feeding in a novel arena after being fasted in their homecage for ∼24 hours (**Figure 6**). The novel arena consisted of the following features: a 40 cm tall white plastic posterboard was wrapped in a circular shape within a 40 cm x 40 cm x 40 cm plastic arena to generate circular arena. Vertical black tape ran down two opposing sides of the circle’s walls. The floor was a perforated metal sheet that had 1 cm wide openings tiled 1 cm apart (edge to edge) throughout the floor, filled in by resting on a piece of opaque plastic. This context was wiped down before/after with 70% EtOH. The food tray was placed next to one of the black stripes. Animals were transported to/from the novel arena in a round white plastic container. To begin, mice were gently placed in the center of the arena.

### Novel open field

Mice were tested for locomotion and anxiety-like behavior in a novel open field (**Figure S10**). The square open field (40 cm x 40 cm x 40 cm) was made of plastic and had solid white walls and a gray floor. Lighting was adjusted to be 100 lux in the center of the arena. Mice were transferred to/from the novel open field in a clear rectangular plastic container. For analyses in Ethovision XT (Noldus), the arena was divided into nine (3 x 3) equally sized squares, with the corner and center squares serving as the “corners” and “center” zones, respectively. To begin, mice were gently placed in the center of the arena.

### c-Fos expression after inhibition of DHPC-DLS in novel homecage

For locomotor tests in a novel clean homecage (**Figure S10**), the chamber was set up identically to tests for spontaneous feeding in the homecage, just without the presence of food or its container. Mice were transferred to/from the clean homecage in a clear rectangular plastic container. Mice started the task by being gently placed in the center of the arena. Locomotion was measured using Ethovision XT (Noldus). 1 hr after the end of exposure to the novel homecage, mice were sacrificed, perfused, and brain tissue processed for c-Fos. Two to three bilateral images of c-Fos in the DLS were generated using am epifluorescence microscope (Nikon) under consistent camera and brightness/contrast settings for all images. Numbers of c-Fos-positive cells (above background) were manually counted per image and averaged per mouse and normalized to counts/mm^2 by converting the quantified pixel area to mm^2 using ImageJ (NIH). Experimenter(s) were blind to group assignments until after counting the images.

### Documentation of optic fiber tracts and viral spread

For optogenetic experiments, experimenter(s) determined the placement of optic fibers based on the path and termination point of tissue damage from the optic fibers in sectioned tissue. Mice were included in the final data set when virus was bilaterally expressed in the target region(s) and optic fibers terminated within 0.5 mm of the target region. The approximate locations of these optic fibers are then visually documented for all mice across all experiments in **Figure S9**. For experiments involving Cre-dependent deletion of *Pdyn* in the septum, we visually documented in the figure the spread of Cre-mCherry expression in the septum and its neighboring regions along multiple points of the anterior-posterior extent of the LS for each mouse. The extent of this spread was then overlayed over each other for each mouse and then then opacity of each was adjusted to 10%, with the scale bar for color intensity noting the level of overlap across the mice. Brain atlas reference images^138^ were digitally modified to remove text and used in figures for schematics or documenting fibers or spread.

### Data exclusions

For circuit mapping experiments, multiple injections across multiple cohorts may have been performed until three to four mice (for numerical quantifications) had minimal or no off-target expression of virus/tracer. Some mapping experiments use a single representative mouse with minimal or no off-target expression of virus/tracer for showing qualitative/representative images (no numerical quantification). Behavioral experiments were performed across one to three cohorts per circuit manipulation (encompassing all groups, as described above). As such, exclusions from any figure are as follows: no mice were excluded from **Figure 1** or **Figure S1**, however it was determined that one mouse in the *Penk* vs. *Pdyn* experiment lacked a collected section of the lateral septum at ∼+0.4 mm from bregma, so only two mice are included in the quantifications for that site. Averages across sites leave out this section for that mouse; its averages are based on its other sections across the LS. No mice were excluded from **Figure S2**. Exclusions are not applicable to **Figure S3**. In **Figure 2**, 9 injected mice were excluded or not fully quantified for having off-target or null viral/tracer expression. In **Figure 3**, 4 injected mice were excluded or not fully quantified for having off-target or null viral/tracer expression. No mice or recorded cells were excluded from the *ex vivo* electrophysiological experiments. In **Figure S4**, 8 injected mice across the experiments were excluded or not fully quantified for having off-target or null viral/tracer expression. In **Figure S5A-S5C**, 4 injected mice across the experiments were excluded or not fully quantified for having off-target or null viral/tracer expression. No mice were excluded from **Figure S5D-S5G**. In **Figure 4**, 1 mouse across the experiments was excluded for having off-target placement of the optic fibers or null expression of the virus. In **Figure S6**, no mice were excluded. For **Figure S7**, 1 mouse was excluded for having off-target placement of the optic fibers or null expression of the virus. For **Figure S8**, any exclusions match those noted for the other figures involving inhibition of DLS(*Pdyn*) cells, DLS(*Pdyn*)-LHA terminals, or DHPC-DLS terminals. **Figure S9** reflects any exclusions noted here. In **Figure 5**, no mice were excluded. In **Figure 6**, no mice were excluded. **Figure S10** reflects any exclusions noted for the other figures that are from DLS(*Pdyn*)-LHA or DHPC-DLS experiments.

### Sex as a biological variable

We did not make any *a priori* predictions for sex differences. Male and female mice were included in most of the experiments in this study. Attempts were made to balance sex within each experiment and group, but we acknowledge that the ratio of males and females are not consistent across every group, and that group sizes may be underpowered for statistical comparisons of sex. Nevertheless, we report in the figures the number of males and females used in each experiment.

### Statistical analyses

Group sizes were based on, and similar to, prior work from our lab exploring DLS circuits and related behaviors^36,50^. For comparisons of means of two groups, parametric two-tailed t-tests were used (paired or unpaired, with or without Welch’s correction, as appropriate). Analysis of variance (ANOVA; factorial or repeated measures, with or without Welch’s correction, as appropriate) was used for comparisons of three or more means or repeated means of three or more or from multiple groups. Geisser-Greenhouse correction was used for ANOVAs of three or more repeated measures. Alpha was set to 0.05 across these tests. To account for multiple comparisons, Tukey’s test was used for post-hoc comparisons of three groups or means, and Bonferroni’s test was used for post-hoc comparisons when four or more means were present. For correlations, simple linear regression was used and R^2 reported. Statistical tests were performed using Prism 10 (GraphPad) and Illustrator 2024 (Adobe) was used for generation of the figures. For simplicity, not all significant post-hoc comparisons (such as following main effects of time) are shown in the figures. Detailed statistical analyses can be found in **Table S1**.

## AUTHOR CONTRIBUTIONS (based on: https://credit.niso.org)

Conceptualization: A.S.; Data Curation: T.D.G., J.B.A., A.B., D.P., M.D.K., A.C.; Formal Analysis: T.D.G., J.B.A., A.B., D.P., M.D.K., A.C.; Funding Acquisition: T.D.G., A.S.; Investigation: T.D.G., J.B.A., A.B., D.P., M.D.K., A.C.; Methodology: T.D.G., J.B.A., A.B., D.P., M.D.K., A.C., X.D., A.S.; Project Administration: A.S.; Resources: X.D., A.S.; Software: T.D.G., J.B.A., A.B., D.P., M.D.K., A.C.; Supervision: A.S.; Validation: T.D.G., J.B.A., A.B., D.P., M.D.K., A.C.; Visualization: T.D.G., J.B.A., A.B., D.P., M.D.K., A.C.; Writing – original draft: T.D.G., A.S.; Writing – review & editing: T.D.G., A.S.

## ACKNOWLEDGEMENTS

We thank Dr. Xiangmin Xu for providing viruses. We thank members of the Sahay lab, L.B.B.S, L.M.S.S, and L.B.G. for comments on the manuscript.

## DATA AND MATERIALS AVAILABILITY

All data in this study are available to any researcher for purposes of reproducing or extending the analyses.

## SUPPLEMENTAL INFORMATION

**Figure S1.**
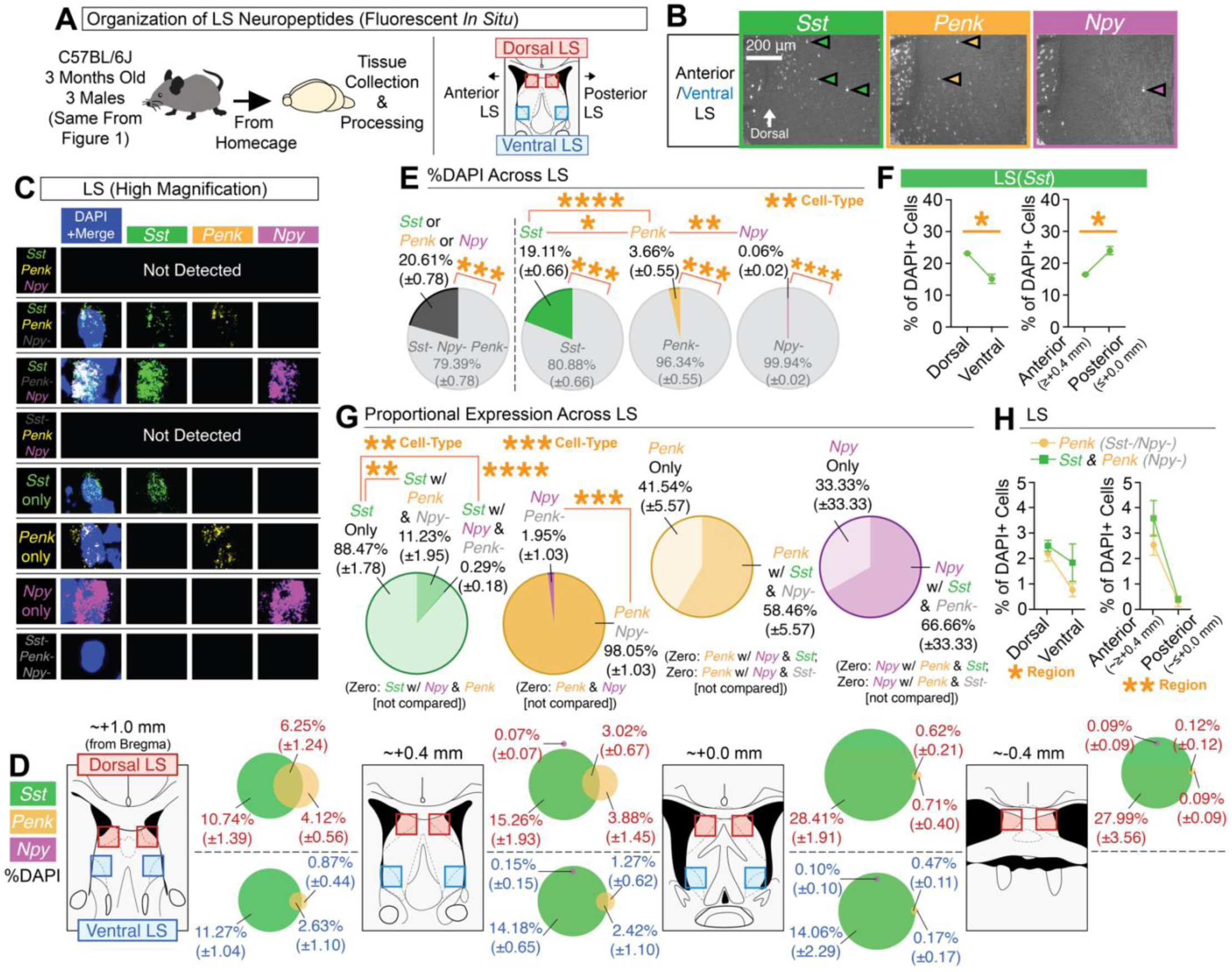
(Related to Main Figure 1). Topographic mapping of neuropeptides reveals partial overlap of *Penk* and *Sst*, whereas *Npy* is sparsely expressed in the LS. **(A)** Multiplex fluorescent *in situ* hybridization was used to map the neuropeptides *Sst*, *Penk*, and *Npy* in the LS across its dorsal-ventral and anterior-posterior regions. **(B)** Representative coronal images of *in situs* for *Sst*, *Penk*, and *Npy* at different regions of the LS. **(C)** High magnification representative images of individual cells for each of the different cell-types detected for *Sst*, *Penk*, and/or *Npy*. **(D)** Mouse atlas images and venn diagrams [%DAPI (±SEM)] depicting the extent of overlap of *Sst*, *Penk*, and/or *Npy* at each quantified region in the LS. **(E)** Average expression (%DAPI) of *Sst*, *Penk*, and/or *Npy* across all quantified regions of the LS. Main effect of cell-type (ANOVA; significant Tukey’s post-hocs) for comparisons of *Sst*-positive, *Penk*-positive, and *Npy*-positive cells. Significant paired t-tests denote comparisons of positive vs. negative expression for each cell-type. **(F)** Average expression (%DAPI) of *Sst* in the dorsal vs. ventral LS and anterior vs. posterior LS (significant paired t-test). **(G)** The average proportion (derived from %DAPI) of each subtype across all quantified LS regions for all *Sst*, *Penk*, and/or *Npy* cells (ANOVA: main effect of cell-type; significant Tukey’s post-hocs), all Penk or Npy cells (significant paired t-test), all *Penk* cells with and without *Sst*, and all *Npy* cells with and without *Sst*. **(H)** Comparisons of the average expression (%DAPI) of *Penk* cells with and without *Sst* in the dorsal vs. ventral and anterior vs. posterior regions of the LS (ANOVAs: main effects for region). For the entire figure, all data are shown as mean (±SEM), and for all statistics: *=p<0.05; **p<0.005, ***p<0.0005; ****p<0.00005.

**Figure S2.**
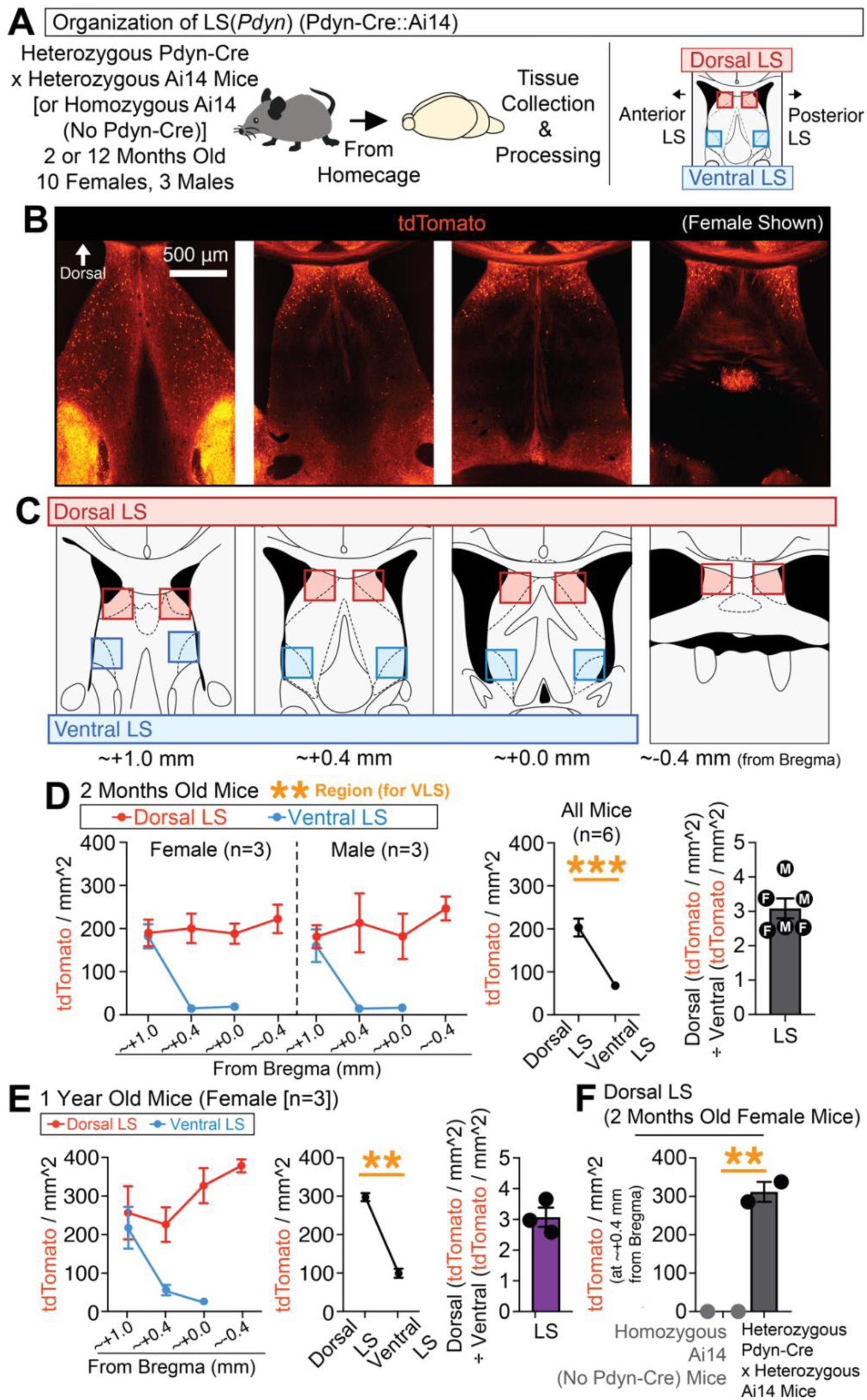
(Related to Main Figure 1). Dorsal bias of developmentally tagged *Pdyn*-expressing cells in the lateral septum of male and female mice. **(A)** Pdyn-Cre mice were crossed with the Cre-dependent tdTomato-expressing Ai14 reporter line to label and quantify *Pdyn*-expressing cells throughout development in their progeny. **(B)** Representative coronal images of tdTomato-expressing cells in the septum of a Pdyn-Cre::Ai14 mouse. **(C)** Mouse atlas images for regions quantified for tdTomato-labeled cells across the LS. **(D)** Quantification of the number of tdTomato cells (per mm^2) per LS region for 2 months old males and females (left panel), as well as the average number of tdTomato cells across the dorsal vs. ventral regions of the LS (middle panel; significant paired t-test). The rightmost graph depicts the average number of dorsal cells divided by the number of ventral cells for all the mice. **(E)** In female mice aged to 1 year, tdTomato cells were quantified across LS subregions (left panel) and compared for average expression across its dorsal and ventral regions (middle panel; significant paired t-test). The rightmost graph depicts the average number of dorsal cells divided by the number of ventral cells for all the mice. **(F)** Comparisons of the number of tdTomato cells in the posterior-dorsal LS in Ai14-positive (Pdyn-Cre-negative) vs. Pdyn-Cre::Ai14 crosses (significant unpaired t-test). For the entire figure, all data are shown as mean (±SEM), and for all statistics: *=p<0.05; **p<0.005, ***p<0.0005; ****p<0.00005.

**Figure S3.**
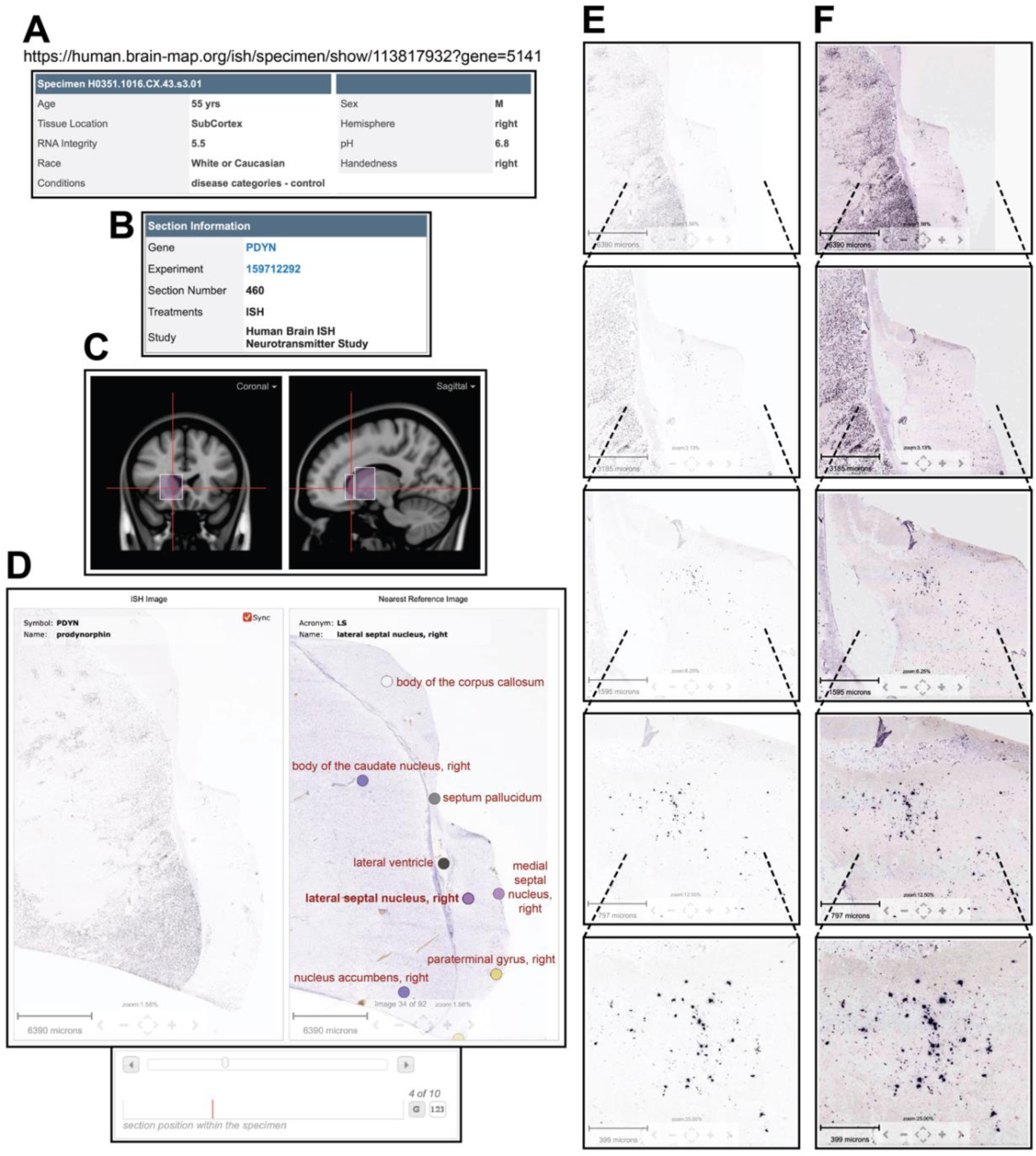
(Related to Main Figure 1). Evidence for conservation of *Pdyn*-expressing cells in the human dorsolateral septum. **(A)** The website link used and a screenshot of the specimen info for *Pdyn* expression in human brain tissue, accessed from the publicly available Human Brain ISH Neurotransmitter Study from the Allen Brain Institute. All images in this figure are screenshots captured directly from this website. **(B)** Screenshot of section information for tissue shown. **(C)** Screenshot of the approximate coronal/sagittal location of tissue shown. **(D)** Screenshots of an *in situ* for *Pdyn* in the human septum and neighboring structures (left panel), with the labeling of the brain structures noted in the Nissl-stained reference image (right panel). The specific image number in the series of *in situ* images is noted in the screenshot in the bottom panel. **(E)** Screenshots of increasing zoom factor (scale bars included) showing Pdyn expression in the human dorsolateral septum of the same section shown in **(D)**. **(F)** The same images from **(E)** but with the contrast artificially enhanced.

**Figure S4.**
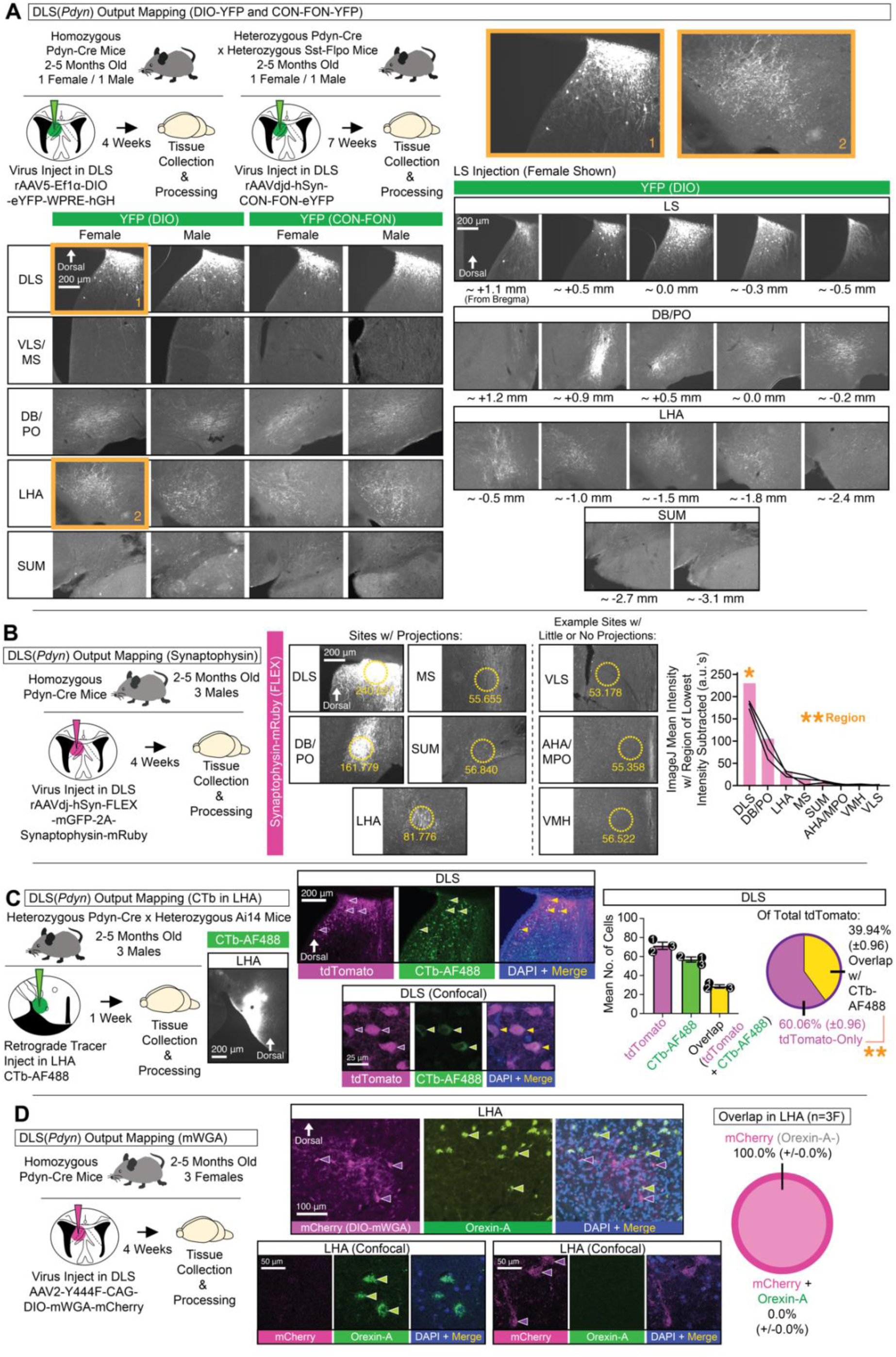
(Related to Main Figure 3). Additional characterization of DLS(*Pdyn*) outputs. **(A)** Pdyn-Cre or Pdyn-Cre::Sst-Flpo mice were injected in the DLS with Cre-dependent or Cre- and Flpo-dependent (CON-FON) YFP-expressing viruses, respectively. Representative coronal images depict YFP expression in DLS(*Pdyn*) cells and their efferent targets in male and female mice. **(B)** Pdyn-Cre mice were injected in the DLS with Synaptophysin-mRuby-expressing virus. Representative coronal images depict Synaptophysin-mRuby expression in DLS(*Pdyn*) cells and their efferent targets and neighboring targets with little or no expression. Dotted-line circles depict example sites and values for measures of intensity in ImageJ (NIH). Rightmost graph plots mean intensity values for each mouse, with the value from the region of lowest intensity subtracted (ANOVA, main effect of region; significant Bonferroni post-hocs). **(C)** Pdyn-Cre::Ai14 mice were injected in the LHA with the retrograde tracer, CTb-AF488. Representative coronal images show expression of tdTomato and CTb-488 in the DLS of Pdyn-Cre::Ai14 mice. Representative confocal image noting overlap of tdTomato with CTb-488 in the DLS. Rightmost graphs show quantifications of the number of tdTomato- and CTb-AF488-expressing cells (no statistical tests used) and their extent of overlap (pie chart: significant paired t-test). **(D)** Pdyn-Cre mice were injected in the DLS with Cre-dependent anterograde virus (DIO-mWGA-mCherry). Representative coronal images show expression of mCherry and Orexin-A in the LHA. Representative confocal images non-overlapping expression of mCherry and Orexin-A, with the rightmost graph showing quantification of this overlap (pie chart: no statistical tests used).

**Figure S5.**
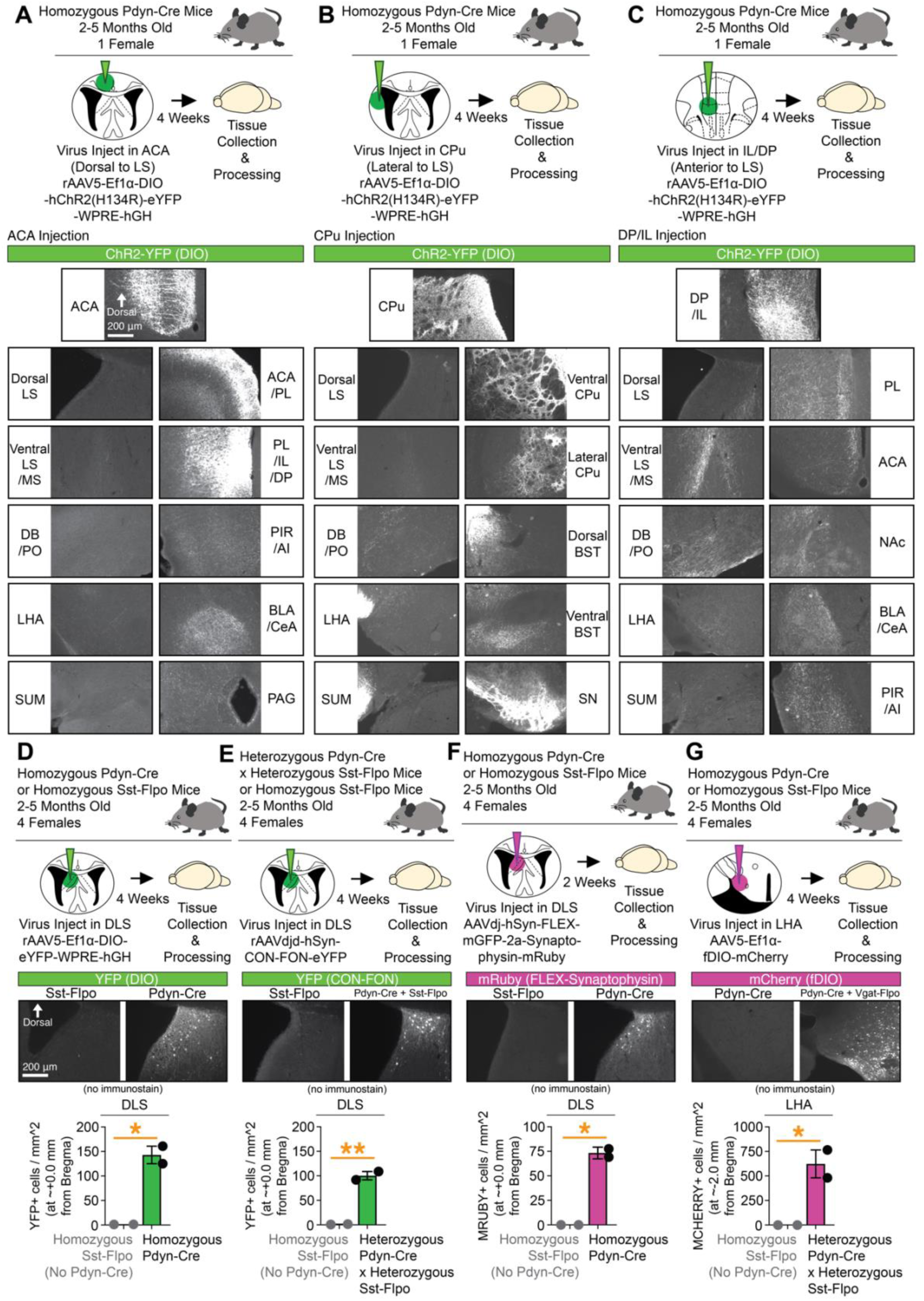
(Related to Main Figure 3). Outputs of DLS-adjacent *Pdyn*-expressing cells and verification of genetic-specificity of viruses. **(A)** Cre-dependent ChR2-YFP-expressing virus was injected into the ACA directly above the DLS in a Pdyn-Cre mouse. Representative images of the injection site, and example sites of output fibers observed are shown (righthand images), along with sites we previously found DLS(*Pdyn*) projections (lefthand images). **(B)** Cre-dependent ChR2-YFP-expressing virus was injected into the CPu directly lateral to the DLS in a Pdyn-Cre mouse. Representative images of the injection site, and example sites of output fibers observed are shown (righthand images), along with sites we previously found DLS(*Pdyn*) projections (lefthand images). **(C)** Cre-dependent ChR2-YFP-expressing virus was injected into the DP/IL directly anterior to the DLS in a Pdyn-Cre mouse. Representative images of the injection site, and example sites of output fibers observed are shown (righthand images), along with sites we previously found DLS(*Pdyn*) projections (lefthand images). **(D)** Specificity of DIO-YFP expression in the DLS was compared in Sst-Flpo vs. Pdyn-Cre mice (significant unpaired t-test). **(E)** Specificity of CON-FON-YFP expression in the DLS was compared in Sst-Flpo vs. Pdyn-Cre::Sst-Flpo mice (significant unpaired t-test). **(F)** Specificity of FLEX-Synaptophysin-mRuby expression in the DLS was compared in Sst-Flpo vs. Pdyn-Cre mice (significant unpaired t-test). **(G)** Specificity of fDIO-mCherry expression in the LHA was compared in Pdyn-Cre vs. Pdyn-Cre::Vgat-Flpo mice (significant unpaired t-test).

**Figure S6.**
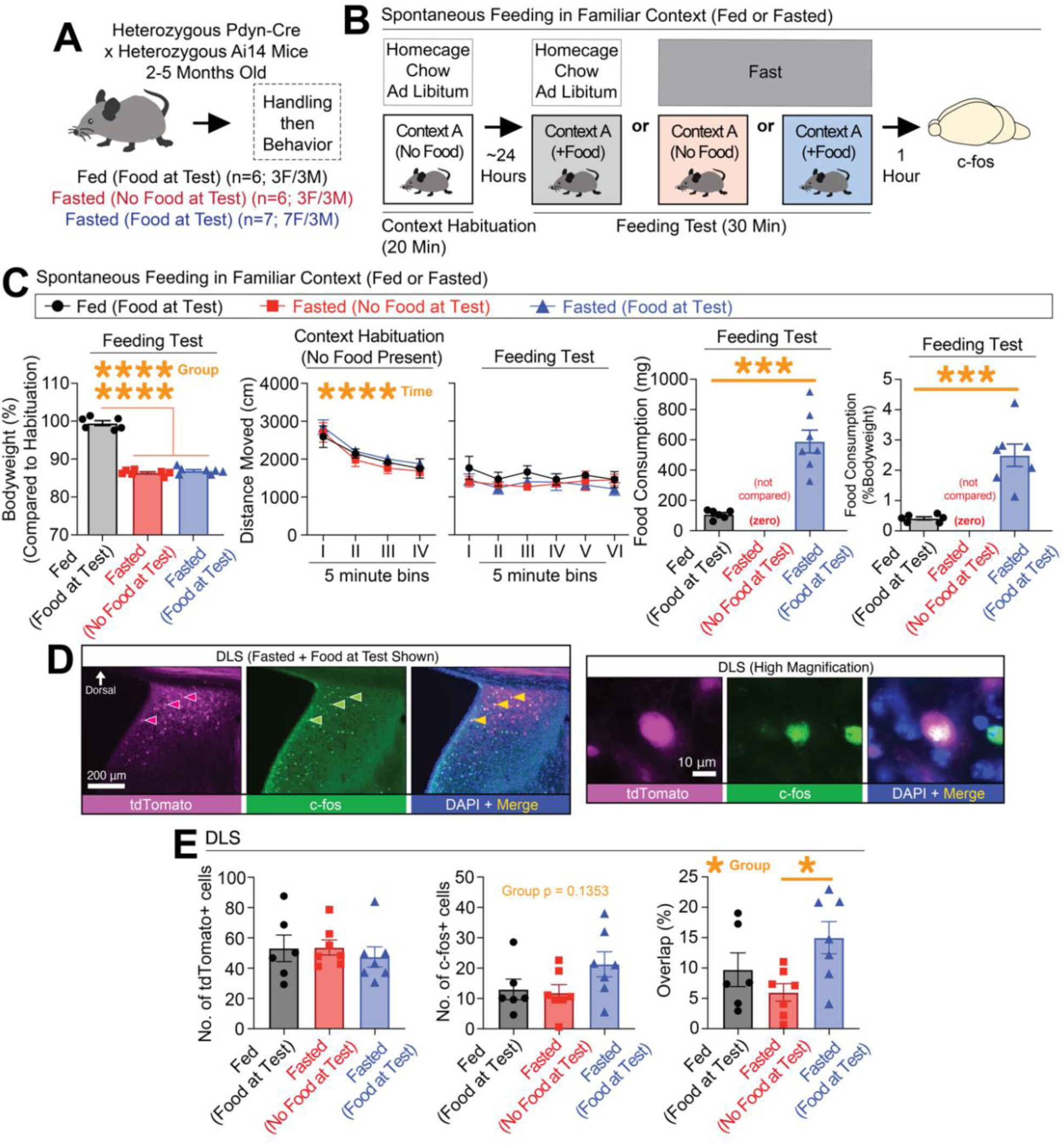
(Related to Main Figure 4). Expression of the immediate early gene, c-Fos, was increased in Pdyn-labeled cells in fasted mice that had access to chocolate flavored food. **(A)** Pdyn-Cre::Ai14 mice were split into three groups. (**B**) Schematic of the behavioral paradigm in which mice were habituated to a context, then either remained on homecage chow or were fasted then given access (or not) to chocolate flavored food in the habituated context. Mice were then sacrificed 1 hour after and processed for c-Fos in the DLS. (**C**) Bodyweight percentages at test, distance moved during each phase (cm), and consumption (mg and %Bodyweight) at test. (**D**) Representative coronal and high-magnification images of c-Fos in the DLS Pdyn-Cre::Ai14 mice that fasted and given food at test. (**E**) Numbers of tdTomato-, c-Fos-expressing, and their overlap (ANOVA, main effect of group, followed by significant Tukey’s post-hocs) in the DLS across groups. For the entire figure, all data are shown as mean (±SEM), and for all statistics: *=p<0.05; **p<0.005, ***p<0.0005; ****p<0.00005.

**Figure S7.**
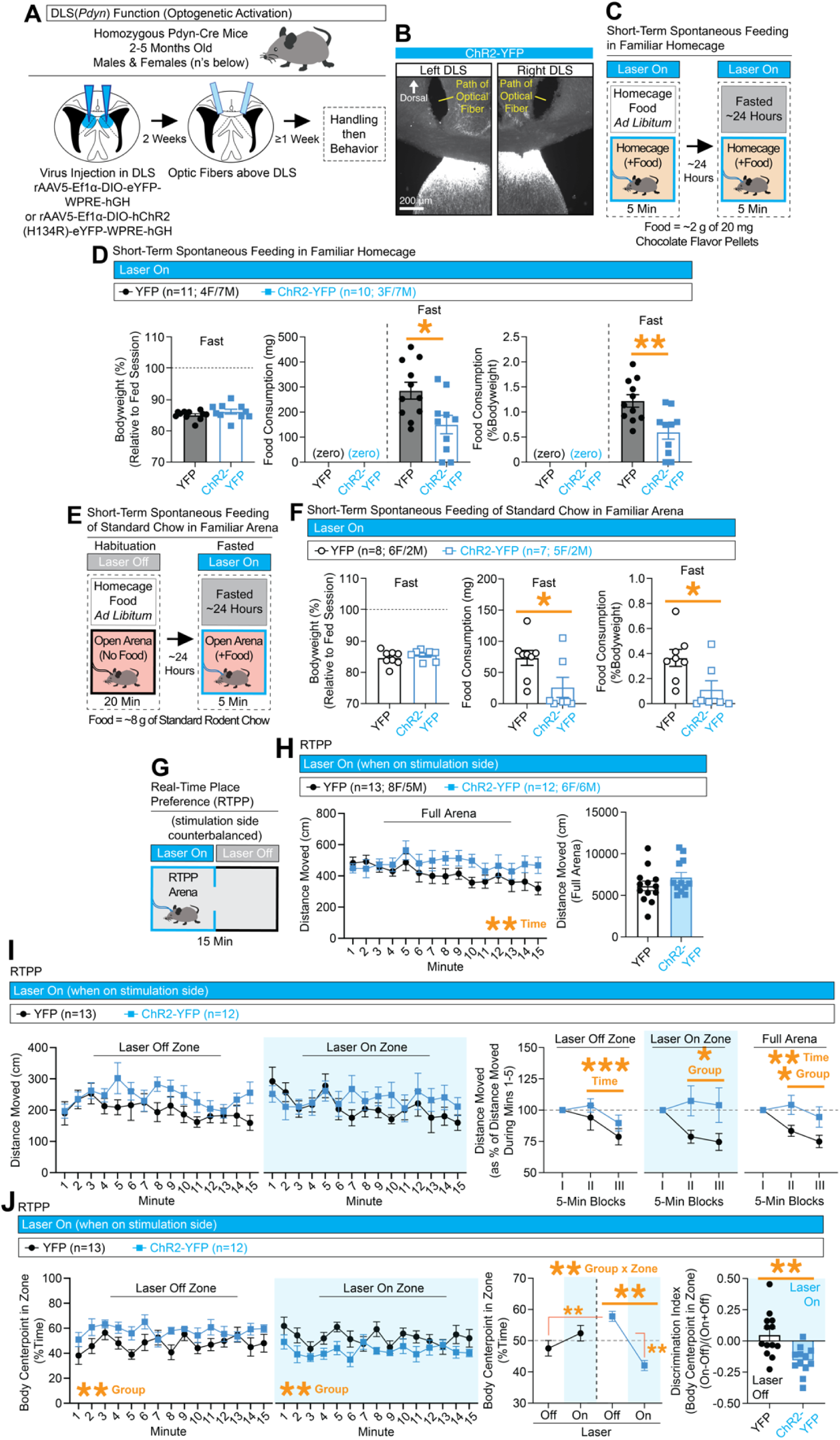
(Related to Main Figure 4). Artificial activation of DLS(*Pdyn*) cells reduces feeding and triggers avoidance. **(A)** Pdyn-Cre mice were injected with Cre-dependent ChR2-expressing or control virus in the DLS and optic fibers were placed above the DLS. **(B)** Representative coronal images with ChR2-YFP expression in the DLS and optic tracts above the DLS. **(C)** Behavioral design for tests for short-term state-dependent spontaneous feeding in the homecage with optogenetic activation. **(D)** Left graph shows bodyweight (%) relative to the first feeding session. Middle and rightmost graphs show the amount of food consumed (in mg and %Bodyweight) during the *ad libitum* and fasted sessions (significant unpaired t-tests for mg or %Bodyweight during fasted session). **(E)** Behavioral design for testing for short-term spontaneous feeding of regular chow in fasted mice in a familiar arena with optogenetic activation. **(F)** Left graph shows bodyweight (%) at the time of the fasted test relative to the day before. Middle and rightmost graphs show the amount of regular chow consumed (in mg and %Bodyweight) during the fasted test (significant unpaired t-tests for mg or %Bodyweight). **(G)** Behavioral design for real-time place preference (RTPP) with optogenetic activation of DLS(*Pdyn*) cells. **(H)** Distance moved at each minute (left graph; ANOVA, main effect of time) and total distance moved (right) during the full test. (I) Leeft graphs show the distance moved per minute in the laser off and on zones. The final graphs show distance moved as a % of the first 5-min block in the laser off zone (repeated measure ANOVA, main effect of time), in the laser on zone (ANOVA, main effect of group), and for the full arena (ANOVA, main effects of time and group). **(J)** Left graphs show %Time per min in the laser off (ANOVA, main effect of group) and laser on zones (ANOVA, main effect of group). Middle graphs show mean %Time in each zone per group across the test (ANOVA, significant zone x group interaction, significant Bonferroni post-hoc for comparing off vs. on zones in ChR2 mice). A discrimination index was generated based on zone preference at test (significant unpaired t-test). For the entire figure, all data are shown as mean (±SEM), and for all statistics: *=p<0.05; **p<0.005, ***p<0.0005; ****p<0.00005.

**Figure S8.**
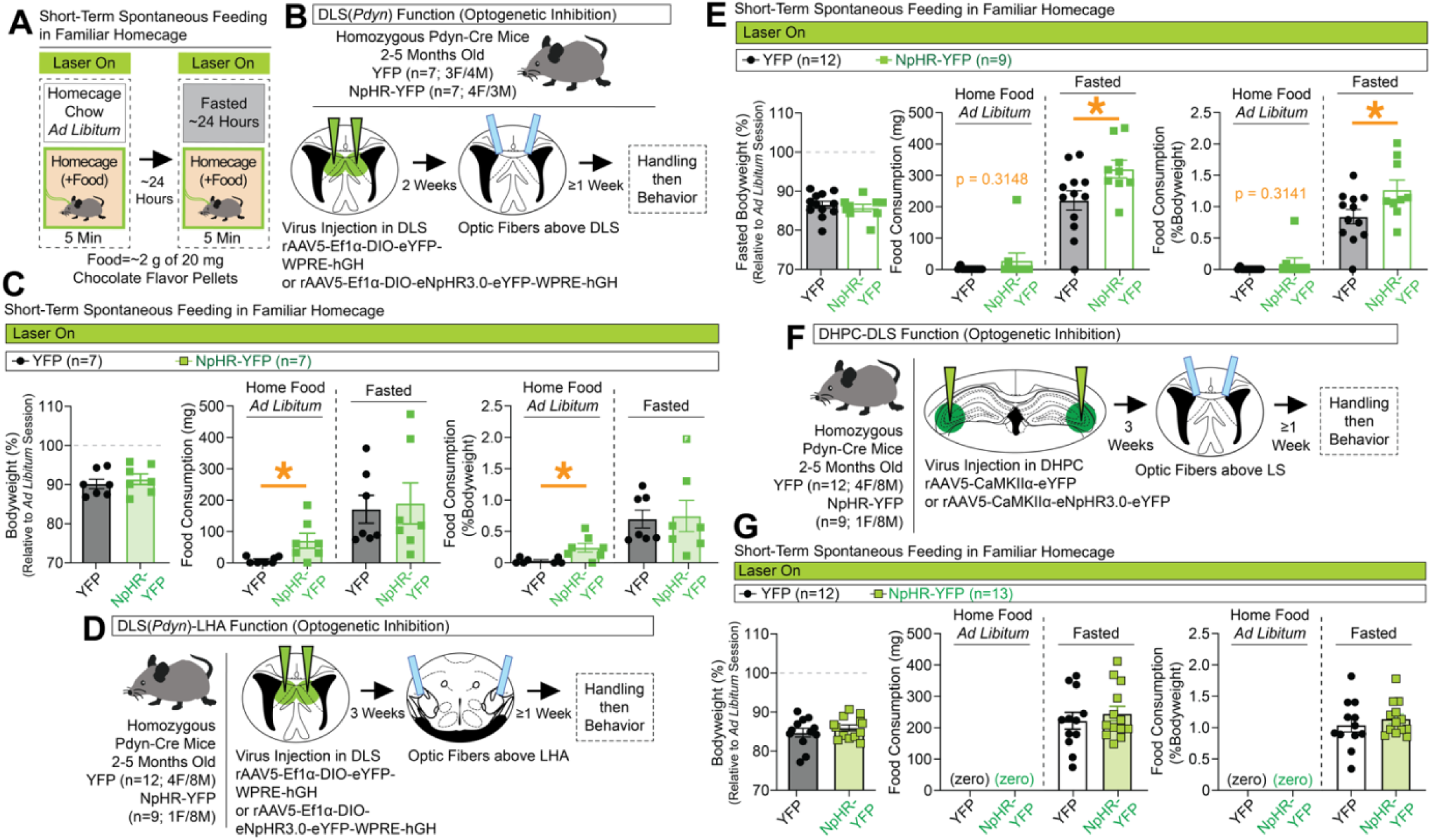
(Related to Main Figure 4). Increases in short-term spontaneous homecage feeding with inhibition of either DLS(*Pdyn*) cells or DLS(*Pdyn*)-LHA terminals, but no changes with DHPC-DLS inhibition. **(A)** Behavioral design for tests for short-term state-dependent spontaneous feeding in the homecage with optogenetic inhibition. **(B)** Pdyn-Cre mice were injected with Cre-dependent NpHR-expressing or control virus in the DLS and optic fibers were placed above the DLS. **(C)** Left graph shows bodyweight (%) relative to the first feeding session. Middle and rightmost graphs show the amount of food consumed (in mg and %Bodyweight) during the *ad libitum* and fasted sessions (significant unpaired t-tests for mg or %Bodyweight during *ad libitum* session). **(D)** Pdyn-Cre mice were injected with Cre-dependent NpHR-expressing or control virus in the DLS and optic fibers were placed above the LHA. **(E)** Left graph shows bodyweight (%) at the time of the fasted test relative to the day before. Middle and rightmost graphs show the amount of food consumed (in mg and %Bodyweight) during the ad libitum and fasted sessions (significant unpaired t-tests for mg or %Bodyweight during the fasted session). **(F)** Pdyn-Cre mice were injected with NpHR-expressing or control virus in the DHPC and optic fibers were placed above the DLS. **(G)** Left graph shows bodyweight (%) relative to the first feeding session. Middle and rightmost graphs show the amount of food consumed (in mg and %Bodyweight) during the *ad libitum* and fasted sessions. For the entire figure, all data are shown as mean (±SEM), and for all statistics: *=p<0.05; **p<0.005, ***p<0.0005; ****p<0.00005.

**Figure S9.**
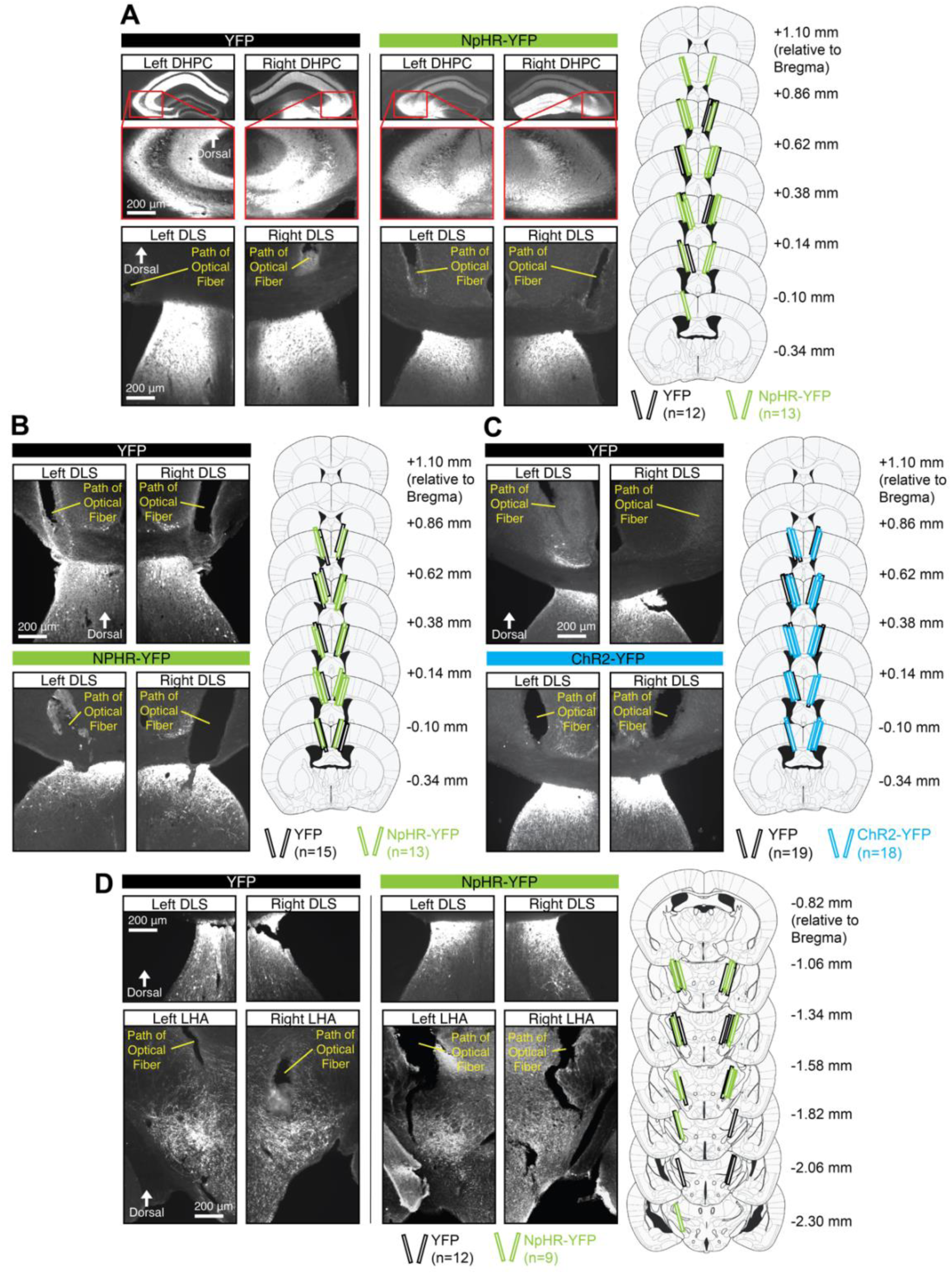
(Related to Main Figure 4). Representative images of viral expression and documentation of placements for optic fibers for behavioral experiments. **(A)** Representative coronal images of bilateral DHPC and DLS showing viral expression of NpHR-YFP or control virus, as well as optic fiber tracts above the DLS. Modified brain atlas images document placements of optic fibers for each group across all DHPC-DLS optical inhibition experiments. **(B)** Representative coronal images of bilateral DLS showing viral expression of NpHR-YFP or control virus, as well as optic fiber tracts above the DLS. Modified atlas images document placements of optic fibers for each group across all DLS(*Pdyn*) optical inhibition experiments. **(C)** Representative coronal images of bilateral DLS showing viral expression of ChR2-YFP or control virus, as well as optic fiber tracts above the DLS. Modified atlas images document placements of optic fibers for each group across all DLS(*Pdyn*) optical excitation experiments. **(D)** Representative coronal images of bilateral DLS and LHA showing viral expression of NpHR-YFP or control virus, as well as optic fiber tracts above the LHA. Modified atlas images document placements of optic fibers for each group across all DLS(*Pdyn*)-LHA optical inhibition experiments.

**Figure S10.**
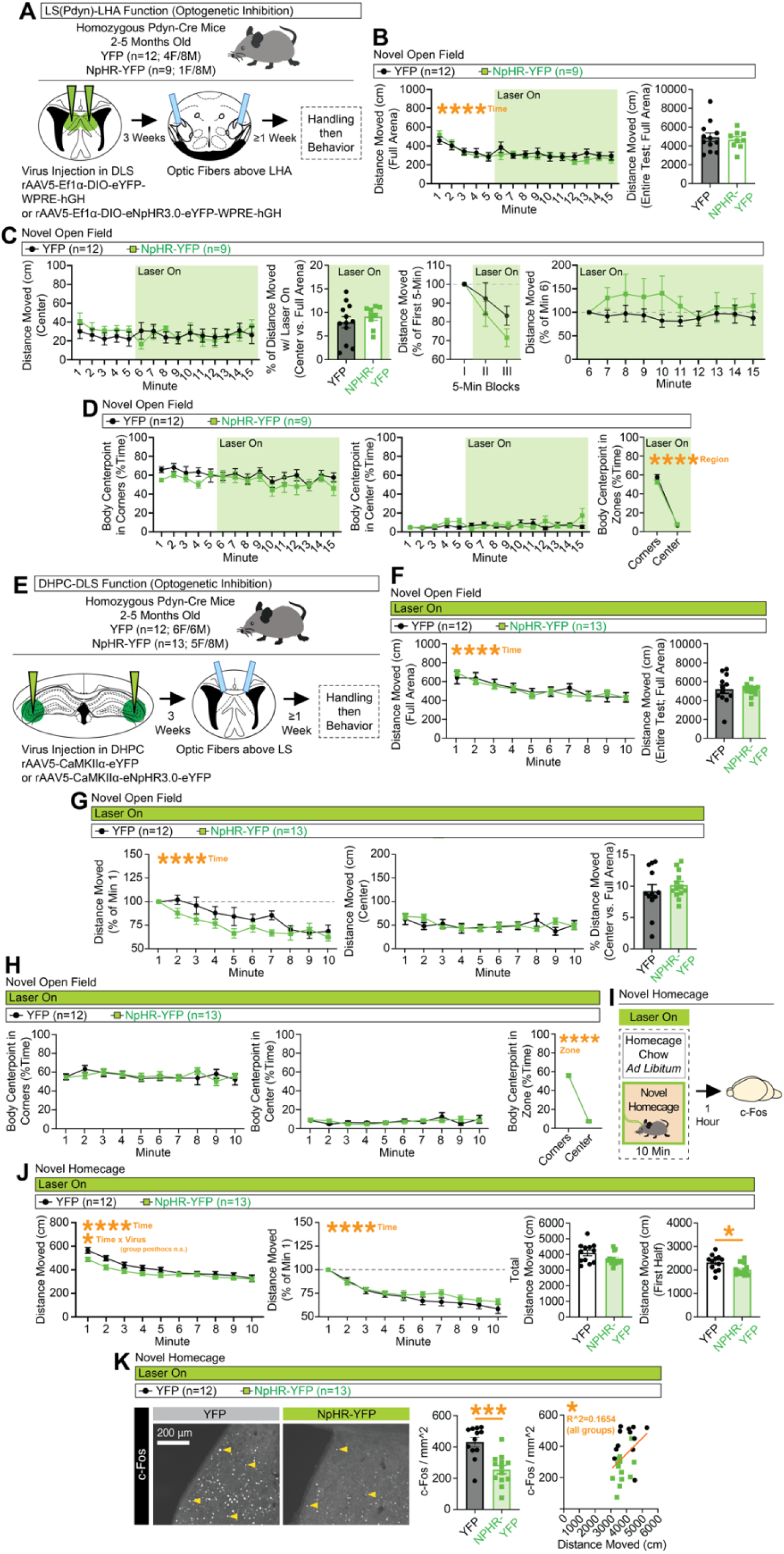
(Related to Main Figure 6). Minimal changes in locomotion or anxiety-like behaviors with optogenetic inhibition of DLS(*Pdyn*)-LHA terminals or DHPC-DLS terminals. **(A)** Pdyn-Cre mice were injected with Cre-dependent NpHR-expressing or control virus in the DLS and optic fibers were placed above the LHA. Mice were tested for locomotion in a novel open field (15 min) with onset of optical inhibition after 5 min. **(B)** Distance moved (cm) in the full arena of the novel open field during each minute (left graph; ANOVA, main effect of time) and total distance moved at test (right graph). **(C)** Distance moved (cm) in the center of the open field during each minute (left graph). Next graph shows the % of total distance that occurred in the center of the arena, followed by a graph showing distance moved in 5-min blocks as a percent of the first 5-min block. The final graph shows distance moved as a percent of movement during minute 6 for each subsequent minute. **(D)** The percent of time the animal spent in any given corner (left graph) or center (middle graph) of the arena for each minute of the test. Final graph compares time spent in the corners vs. center for full duration of the laser on (ANOVA, main effect of zone). **(E)** Pdyn-Cre mice were injected with NpHR-expressing or control virus in the DHPC and optic fibers were placed above the DLS. Mice were tested for locomotion in an open field and novel homecage. **(F)** Distance moved (cm) in the full arena of the novel open field during each minute (left graph) and total distance moved at test (right graph). **(G)** Distance moved as a % of the first minute for each minute of the test (left graph; ANOVA, main effect of time). Middle graph shows distance moved (cm) in the center of open field. Final graph shows the % of total distance that occurred in the center of the arena. **(H)** The % of time the animal spent in any given corner (left graph) or center (middle graph) of the arena for each minute of the test. Final graph compares time spent in the corners vs. center for the entire test (ANOVA, main effect of zone). **(I)** These same DHPC-DLS mice were exposed to a novel homecage before being sacrificed for c-Fos. **(J)** Distance moved (cm) in the full homecage during each minute (left graph; ANOVA, main effect of time, time x virus interaction). Next graph shows distance moved as a % of the first minute for each minute of the test (ANOVA, main effect of time). Final two graphs show distance moved during the whole test or first 5 min (significant unpaired t-test). (**K**) Representative coronal images showing c-Fos expression in the DLS of NpHR-expressing and control mice (left), this expression quantified (middle; significant unpaired t-test), and the correlation of c-Fos with distance moved across the test (significant R^2, across groups). For the entire figure, all data are shown as mean (±SEM), and for all statistics: *=p<0.05; **p<0.005, ***p<0.0005; ****p<0.00005.

## KEY RESOURCES TABLE

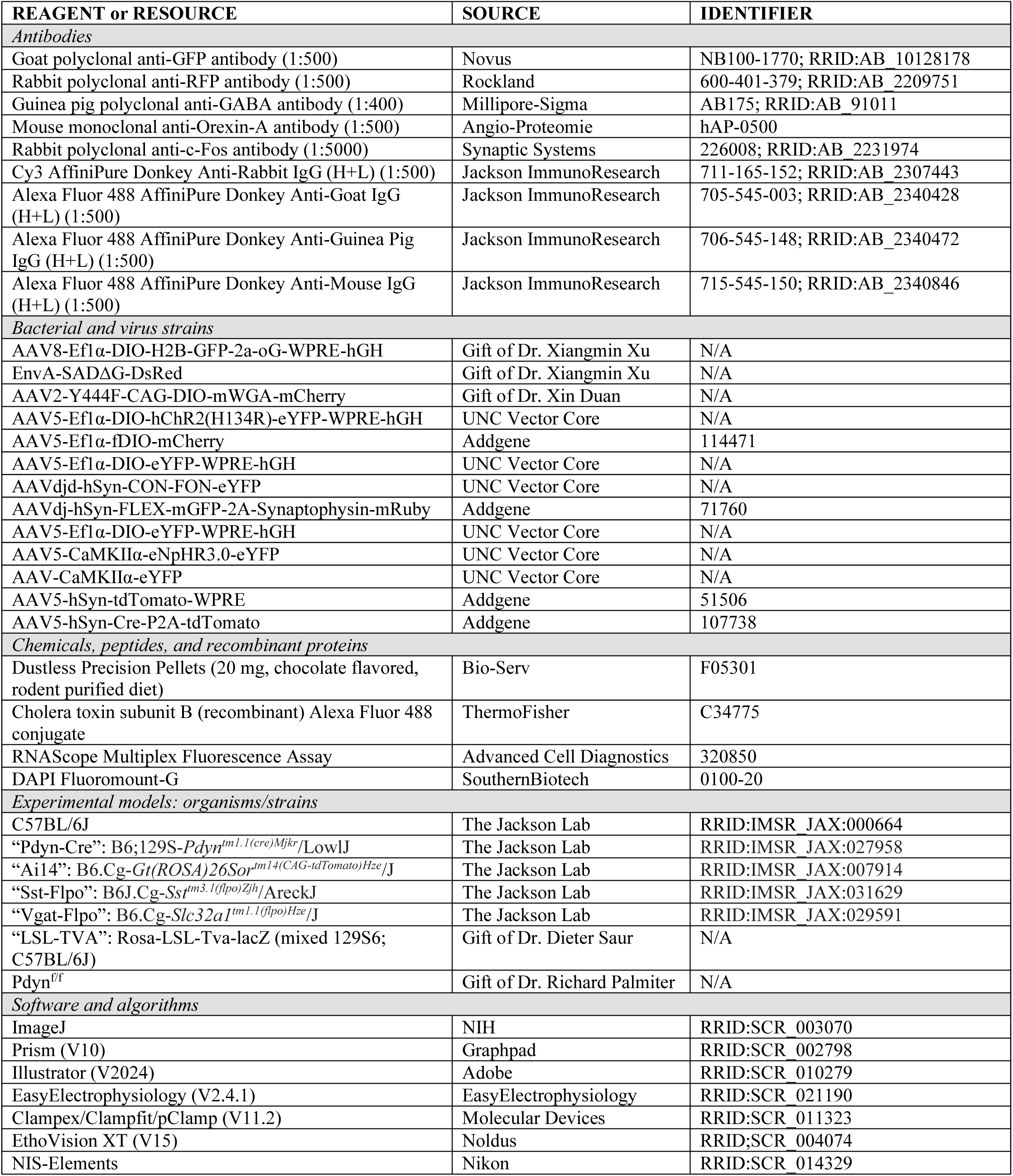

